# Are asymmetric inheritance systems an evolutionary trap? Transitions in the mechanism of genome loss in the scale insect family Eriococcidae

**DOI:** 10.1101/2022.06.23.497384

**Authors:** Christina N Hodson, Alicia Toon, Lyn Cook, Laura Ross

## Abstract

Haplodiploidy and paternal genome elimination (PGE) are examples of asymmetric inheritance, where males transmit only maternally inherited chromosomes to their offspring. Under haplodiploidy this results from males being haploid, whereas under PGE males inherit but subsequently eliminate paternally inherited chromosomes during meiosis. Their evolution involves changes in the mechanisms of meiosis and sex determination, and sometimes also dosage compensation. As a result, these systems are thought to be an evolutionary trap, meaning that once asymmetric chromosome transmission evolves, it is difficult to transition back to typical Mendelian transmission. We assess whether there is evidence for this idea in the scale insect family Eriococcidae, a lineage with PGE and the only clade with a suggestion that asymmetric inheritance has transitioned back to Mendelian inheritance. We conduct a cytological survey of 13 eriococcid species, and a cytological, genetic, and gene expression analysis of species in the genus *Cystococcus*, to investigate whether there is evidence for species in this clade evolving Mendelian chromosome transmission. Although we find that all species we examined exhibit PGE, the mechanism is extremely variable within Eriococcidae. Within *Cystococcus*, in fact, we uncover a previously undiscovered type of PGE in scale insects, where in males paternally inherited chromosomes are present, uncondensed, and expressed in somatic cells, but are eliminated prior to meiosis. Broadly, we fail to find evidence for a reversion from PGE to Mendelian inheritance in Eriococcidae, supporting the idea that asymmetric inheritance systems such as PGE may be an evolutionary trap.

## Introduction

Reproduction systems (i.e., chromosome inheritance and sex determination) are extraordinarily variable across the tree of life. Understanding the forces that have led to this variety, and what leads to or constrains transitions to new systems has been a major goal in evolutionary biology in recent years (Bachtrog *et al*. 2014). In most species chromosome inheritance is a fair process overall, such that either homologous chromosome in a diploid has an equal probability of being transmitted to offspring (i.e., Mendelian chromosome transmission). However, some reproductive systems are characterised by unequal chromosome transmission (asymmetric inheritance or non-Mendelian transmission): haplodiploidy and paternal genome elimination (PGE) are the two most widespread examples of these (de la Filia *et al*. 2015; Ross *et al*. 2022). In both haplodiploidy and PGE, males transmit only maternally derived chromosomes to their offspring. However, while under haplodiploidy males develop from unfertilized eggs, in PGE males develop from fertilized eggs and are diploid, but paternally inherited chromosomes are not incorporated into viable sperm (Normark 2003). It remains unclear how haplodiploidy and PGE evolve, yet these systems appear to be extraordinary successful and have evolved repeatedly, with more than 20 origins across invertebrates. They are found across large clades, including entire orders (e.g. Hymenoptera, thrips, parasitic lice and globular springtails) and make up around 12% of extant animal species (de la Filia *et al*., 2015).

Ideas about the evolution of haplodiploidy and PGE suggest that genomic conflict, specifically conflict between the maternal and paternal halves of the genome, drive transitions to these systems (Hartl and Brown 1970; Bull 1979, 1983; Burt and Trivers 2006; Gardner and Ross 2014; Ross *et al*. 2019). In these models, asymmetric transmission benefits maternally derived chromosomes/genes in males as they are transmitted to all male offspring (rather than 50%, as expected under Mendelian transmission). Therefore, we might expect the evolution of asymmetric inheritance to be quite dynamic, especially in early stages, with frequent transitions between Mendelian and non-Mendelian inheritance depending on which party in conflict gains the upper hand. An indication of this conflict would be species with Mendelian inheritance within clades with asymmetric inheritance. Currently, there is not a single record of transitions from haplodiploidy back to Mendelian inheritance and the evidence for transitions within PGE clades remains unconfirmed. Transitions from Mendelian to asymmetric inheritance involves a number of changes. Changes in the mechanism of meiosis, sex determination, ploidy, and dosage compensation, among others, often occur with a shift to asymmetric inheritance (Gardner and Ross, 2014; Ross *et al*., 2019). Because of the number and complexity of these changes, asymmetric inheritance systems are thought to be an evolutionary trap (i.e., once they evolve it may be difficult to transition back to Mendelian inheritance) (Bull, 1983; Bachtrog *et al*., 2014). The apparent absence of transitions from haplodiploidy to diplodiploidy—or more precisely a lack of diploid species within haplodiploid clades (Tree of Sex database-http://www.treeofsex.org/)—has been taken as evidence for such an evolutionary trap (Bull 1983; Bachtrog *et al*. 2014; Blackmon *et al*. 2017). It is less clear if PGE is a similar evolutionary trap or if reversions to Mendelian reproduction are possible.

Unlike under haplodiploidy, males in species with PGE are often diploid with paternal chromosomes eliminated only in germline cells undergoing meiosis. So, transitions back to diplodiploidy may not involve a change in male ploidy, and spermatogenesis often still involves two meiotic divisions. However, the evolution of PGE does involve significant changes in the mechanism of meiosis such that chromosomes segregate according to their parent of origin and, in many species with PGE, meiosis is also altered in other ways. For instance, in scale insects, the meiotic sequence is reversed (i.e. inverted meiosis) and in both scale insects and fly lineages with PGE, a highly derived monopolar spindle is present in male meiosis which only attaches to maternally inherited chromosomes (Brown 1967; Kubai 1982; Bongiorni *et al*. 2004). Furthermore, all documented lineages with PGE have evolved from an XO or XY sex determination systems and the evolution of PGE from these sex chromosome systems requires a change in the sex determination mechanism, since otherwise males in PGE systems would always transmit the maternally-derived X chromosome through sperm, resulting in female-only offspring (Gardner and Ross, 2014). Therefore, lineages with PGE have evolved unconventional sex determination systems, either involving elimination of sex chromosomes after fertilization or silencing/elimination of paternally inherited chromosomes in males (Du Bois 1933; Nur 1990; Bongiorni *et al*. 2001). Additionally, although males generally remain diploid, in some PGE systems the mechanism of PGE has evolved such that some paternally inherited chromosomes are eliminated or transcriptionally repressed in cells of males early in development, and therefore the evolution of PGE causes males to exhibit haploid rather than diploid gene expression in somatic cells (Brown and Bennett 1957; de la Filia *et al*. 2021).

So, is there any evidence that Mendelian reproduction has re-evolved within PGE clades? PGE has evolved independently at least seven times, and although it generally occurs across large clades, many of these are poorly studied both in terms of biology and systematics (Gardner and Ross, 2014; de la Filia *et al*., 2015). The only clade for which a considerable number of taxa (around 500 or ∼5% of described species) have been studied are scale insects (Hemiptera: Coccoidea) (Nur 1980 p. 19; Gavrilov 2007; Ross *et al*. 2010). The phylogenetic distribution of PGE across this clade suggests that transitions back to Mendelian chromosome inheritance are rare, but have potentially occurred. In mealybugs, there is evidence that some paternal chromosomes occasionally escape elimination during meiosis (de la Filia *et al*. 2019). Additionally, in two scale insect species, *Stictococcus* sp. and *Lachnodius eucalypti*, cytogenetic analyses suggest that PGE may be absent (Brown 1977; Nur 1980). These two species are relatively closely related, and belong to a scale insect clade with substantial variability in the mechanism of PGE (Eriococcidae *sensu lato*). This variability has been argued to be due to an evolutionary arms race between maternal and paternal alleles over paternal transmission in males, suggesting that scale insects, and in particular, the clades in which *Stictococcus* sp. and *L. eucalypti* belong, should be investigated in more detail, as this may be a group in which conflict over chromosome transmission to future generations is high (Brown 1964; Herrick and Seger 1999; Ross *et al*. 2010).

Here we focus on scale insects within the scale insect family Eriococcidae, and explore whether there is any evidence for paternally inherited chromosomes regaining expression/transmission through males. The Eriococcidae belong to Neococcidae, a large monophyletic clade (14 families, 6000 species) of scale insects that have evolved PGE (Ross *et al*., 2010). Although males across the clade do not transmit paternally inherited chromosomes through sperm, there is variability in the mechanism of PGE between species; both during meiosis and during early embryogenesis (**Figure 1**). The differences in the mechanism of PGE is classified into three categories within scale insects, which differ in whether paternally inherited chromosomes are retained and heterochromatized (epigenetically silenced) in embryogenesis (Leccanoid and Comstockiella systems: **Figure 1**) or whether they are eliminated entirely in embryogenesis (Diaspidid system: **Figure 1**). In both cases, males have no or limited expression of paternal chromosomes in somatic cells (de la Filia *et al*. 2021). Additionally, during meiosis, the three categories differ in whether there is one division in meiosis (Diaspidid/ sometimes Comstockiella) or two (Leccanoid/ sometimes Comstockiella), and whether any paternally inherited chromosomes are eliminated prior to meiosis (Comstockiella) (Nur, 1980; Ross *et al*., 2010).

**Figure 1.**
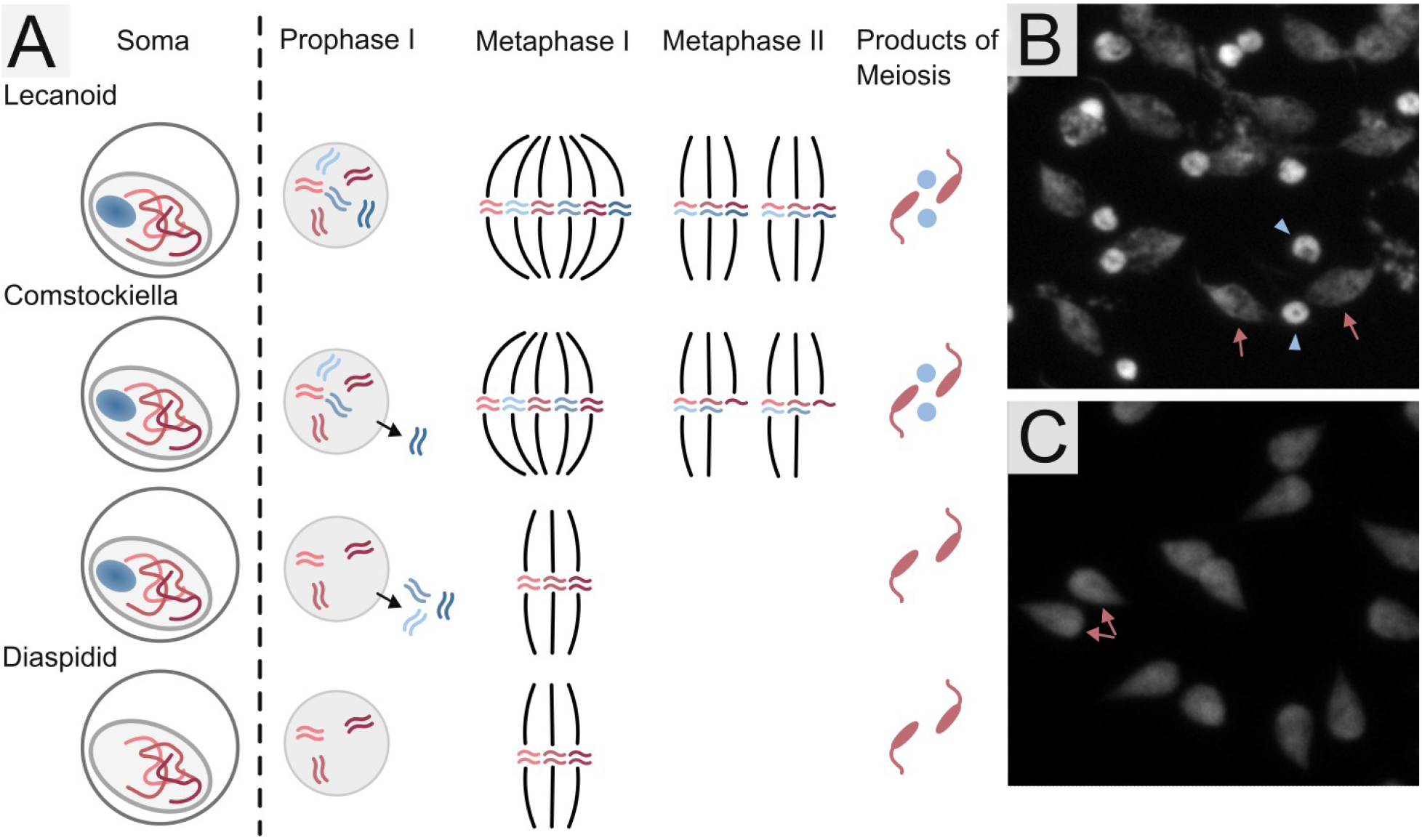
Differences in the mechanism of PGE in scale insect species. Scale insects exhibit inverted meiosis (i.e. sister chromatids separate in the first division of meiosis while homologous chromosomes separate in the second division) and chromosomes segregate according to their parent of origin. In the Lecanoid system, paternally inherited chromosomes (blue, maternal chromosomes in red) are retained in males throughout development, and are condensed as heterochromatic bodies in somatic cells (blue circle representing the condensed paternal chromosomes in a ball). During meiosis, paternal chromosomes are present throughout meiosis, but form into non-viable pycnotic nuclei (blue circles), which are degraded following meiosis. In the Comstockiella system, somatic cells of males look similar to Lecanoid species, but some paternal chromosomes are eliminated from cells just prior to meiosis. The number eliminated is variable between/within species, and pycnotic nuclei can be either present or absent depending on the number eliminated. In the Diaspidid system, paternally inherited chromosomes are eliminated from all cells of males in early development, so heterochromatic bodies are absent from the male soma and pycnotic nuclei are not present after meiosis, which has only one division (as there are no paternal chromosomes). **B** Image of sperm elongating following meiosis in a species with pycnotic nuclei (*Pseudococcus viburni*-image courtesy of Isabelle Vea). **C** Image of sperm elongating following meiosis in a species without pycnotic nuclei (*Cystococcus campanidorsalis*). Blue arrows point to pycnotic nuclei while pink arrows point to elongating sperm.

Eriococcid scale insects exhibit Comstockiella PGE, which is thought to be the ancestral reproduction system in this clade, but there is remarkable diversity in how meiosis occurs in different species (Gavrilov, 2007; Ross *et al*., 2010). For instance, the number of paternal chromosomes eliminated in males prior to meiosis differs between and even within species, leading to differences in the number of divisions in meiosis and whether pycnotic nuclei form after meiosis (Brown, 1967; Nur, 1980). For both of the suggested losses of PGE in Eriococcidae, in *Lachnodius eucalypti* and *Stictococcus* sp., there is no clear evidence that paternal chromosomes are eliminated in male meiosis and - unlike in other scale insects with PGE – somatic cells in males lack heterochromatic bodies containing the silenced paternal genome. This led researchers to suggest that PGE was absent in these species (Brown, 1967; Nur, 1980). However, this conclusion was based on cytological observations of a small number of specimens, and both of these species have uncertain phylogenetic placements, with *L. eucalypti* thought to belong to the Gondwanan clade of eriococcid scale insects, and *Stictococcus* sp. now potentially placed in Stictococcidae which is nested inside Eriococcidae *sensu lato* (Cook *et al*. 2002; Cook and Gullan 2004; Gullan and Cook 2007). Therefore, although Eriococcidae and related species are a promising group for exploring transitions from PGE to Mendelian inheritance, a more thorough examination, encompassing genetic as well as cytological data, is needed to understand whether and how transitions in the reproductive system have occurred, and whether transitions have occurred once, or several times.

We investigate whether there is evidence for transitions from PGE to Mendelian inheritance in species within Eriococcidae using cytological, genetic and transcriptomic analyses. In eriococcidae species that exhibit Comstockiella PGE, somatic cells of males have heterochromatic bodies containing paternal chromosomes. We therefore first examined somatic tissue in 13 Eriococcidae species to determine whether heterochromatic bodies are present in somatic cells of males. We then investigate male meiosis in a subset of species *(Ascelis*/*Cystococcus* clade) which show variation in whether heterochromatic bodies are visible. Further, we study the inheritance patterns (using microsatellite markers) in two species of *Cystococcus*: *C. campanidorsalis*, in which male heterochromatic bodies are present, and *C. echiniformis*, in which male heterochromatic bodies are absent. This allowed us to determine whether inheritance patterns from mothers to sons is consistent with what we would expect under PGE inheritance. Finally, we generate RNAseq data from females and two offspring for *C. echiniformis* and *C. campanidorsalis* families to determine whether a loss of heterochromatic bodies corresponds to a transition in gene expression profiles such that paternally inherited genes are expressed (i.e., diploid rather than haploid expression of chromosomes). We find that although some species within Eriococcidae do not have heterochromatic bodies in somatic tissue in males, there is no evidence that PGE has been lost in these species, rather, there has been a transition to a different type of PGE, unlike those described so far in scale insects. Interestingly, we also find that both *C. campanidorsalis* and *C. echiniformis*, which differ in whether they have heterochromatic bodies, have substantial expression of paternally inherited chromosomes in males. This suggests that this lineage has undergone a change from uniparental to biparental expression in the past and also that the presence of heterochromatic bodies does not always correspond to a complete lack of paternal chromosome expression in males with PGE. More broadly, we find no evidence for a transition from PGE back to Mendelian reproduction in eriococcid scale insects, and thus fail to reject the idea that PGE is an evolutionary trap.

## Materials and Methods

### Study system

Eriococcidae is a diverse and widespread family of scale insects. We focus primarily on the monophyletic subclade labeled the Gondwanan clade by Cook and Gullan (2004), which has approximately 70 species, is relatively well characterized, and contains *Lachnodius eucalypti* (a species reported to lack PGE). All species are ectoparasites of trees and shrubs in the Myrtle family (Myrtaceae). Most species are highly host-plant specific and many in the clade form galls (Cook and Gullan, 2004). Although species within Eriococcidae can vary in chromosome number from 2n = 4-192 (Cook 2000), the chromosome number in most species in the Gondwanan clade is 2n=18 where known (Brown, 1967; Cook, 2000). Because the chromosomes in some species are too numerous and small to count individually, we confined our analyses to the behaviour of the maternally vs. paternally inherited chromosomes in males.

### Staining of male somatic tissue for the presence of heterochromatic bodies

Samples were collected in Queensland and New South Wales, Australia, from 2010 to 2017 (see **Supplementary Table 1** for details) and stored in 1:3 glacial acetic acid: ethanol fixative at 4°C. We performed DAPI staining on somatic tissue of males for 13 species of Eriococcidae to determine whether heterochromatic bodies are present. The majority of species had not previously been studied cytologically, although we also included *Ascelis schraderi* and *Eriococcus coriaceus*, which had previously been examined by Brown (1967), to ensure that our staining techniques were producing comparable results. We did not include any species in the genus *Apiomorpha*, as 45 species within this family have already been examined, all of which were found to have heterochromatic bodies in males (Cook, 2000). With the species examined previously and in our study, we now have information about whether somatic heterochromatization is present in 79 species of Eriococcidae.

Many eriococcid species are sexually dimorphic at an early stage and some exhibit sexual dichronism (i.e. male and female offspring are produced at different times in the female reproductive period) (e.g., *Cystococcus*, Gullan and Cockburn 1986). Thus it is often possible to sex individuals, even at very early stages of development. We preferentially stained males from later life stages but, and when we were unable to determine the sex (embryos or first instar larvae) we stained at least 10 individuals to try to ensure that we stained some males (note that for all samples for which we were unable to determine the sex of individuals, we also stained older males from the same species; **Supplementary Table 2**). Otherwise, we stained 2-4 males from each species at each sampling location by spreading the body tissue of each male thinly on a slide and staining with DAPI (further details in Supplementary Methods).

Heterochromatic bodies appear as a densely stained body at the periphery of the cell nucleus in somatic cells (Bongiorni *et al*., 2001). However, in scale insects with PGE, somatic heterochromatization is not always present in all types of somatic cells (Nur 1967) and in some cases we could not sex individuals before staining. Therefore, we determined whether heterochromatic bodies were present in any somatic cells in each slide and any species in which we were able to identify heterochromatic bodies in somatic tissue of any individual we marked as possessing heterochromatic bodies (see **Supplementary Table 2** for information on number of slides/ stages and sex of specimens stained for each species).

### Staining of male meiosis in *Cystococcus/ Ascelis*

For species within the *Cystococcus/ Ascelis* clade, we also performed DAPI staining on tissue undergoing meiosis in males. We stained males at the late 3^rd^ instar larval/ early pupal development stages, as we found that this is the stage when meiosis occurs in males. The staining procedure was the same as for somatic tissue with the exception that, when we dissected the individual, we removed as much somatic tissue as possible from the preparation. Meiosis in male scale insects takes place in a cyst, in which the number of nuclei per cyst is generally consistent in each species (Brown, 1967). This allowed us to determine which stage of meiosis was occurring by counting the number of nuclei in each sperm cyst. Additionally, if there are two divisions in meiosis and PGE, the final products of meiosis are spermatids containing maternally inherited chromosomes that elongate into functional sperm and pycnotic nuclei, containing paternally inherited chromosomes (**Figure 1**). Therefore, the presence of pycnotic nuclei at the end of meiosis indicates that the species undergoes PGE. We scored slides by evaluating first whether pycnotic nuclei were present after meiosis, then by examining the number of divisions in meiosis. We did this by counting the number of cells in each sperm cyst and the stage of meiosis that was occurring (i.e. prophase, metaphase, anaphase etc.) and counting how many sperm were present in sperm bundles at the end of meiosis. Then we were able to determine how many divisions took place in meiosis by evaluating whether the number of sperm were 4x more than the number of nuclei in primary spermatids (2 divisions in meiosis) or 2x more than the number of nuclei in primary spermatids (1 division in meiosis). Although, note that we were not able to view all stages of meiosis for all the species of interest (**Supplementary Table 2**).

### Eriococcidae phylogeny

We estimated a phylogeny with all species for which we stained male somatic tissue, along with some additional species within Eriococcidae that had previously been examined for somatic heterochromatization (Brown, 1967) (**Supplementary Table 2**), using the mitochondrial CO1 and the nuclear rRNA 18S genes. For both the CO1 and 18S loci, some of the nucleotide sequences we used were from previously published studies (Cook *et al*. 2002; Cook and Gullan 2004; Gullan and Cook 2007; Semple *et al*. 2015) (see **Supplementary Table 3** for accession numbers for published sequences). For the nucleotide sequences that we generated ourselves, we sequenced an approximately 600bp region of the small subunit ribosomal RNA gene 18S. We conducted PCR reactions with the 2880/Br primer set (von Dohlen and Moran 1995) and Sanger sequenced the products in both directions. For the CO1 locus, we used approximately 510bp of the 5’ region (COI barcode region), Sanger sequenced in both directions with either the mitochondrial primers PCO_F1 (Park *et al*. 2010) and HCO (Folmer *et al*. 1994) (for the majority of species) or the primers CystCOIF and CystCOIR (Semple *et al*., 2015) (for *Cystococcus* and *Ascelis* species) (See **Supplementary Table 4** for primer information and thermocycling conditions). We aligned CO1 and 18S sequences separately in Geneious Prime (2020.2.3) with the MAFFT plugin (v 7.450) (Katoh and Standley 2013). For CO1 sequences, we did a translation alignment using ‘invertebrate mitochondrial’ as the genetic code. We then used IQtree to generate maximum likelihood phylogenies with parameters -alrt 1000 -B 1000, which estimates ultrafast bootstrap values and SH-aLRT values at nodes with 1000 replicates each, and allows IQtree to select the most appropriate substitution model (v2.0.3) (Guindon *et* al. 2010; Kalyaanamoorthy *et al*. 2017; Hoang *et al*. 2018; Minh *et al*. 2020). An ultrafast bootstrap value of >= 95% and an SH-aLRT value of >= 80% indicates high confidence in that clade given the data. We specified the outgroup as *Parasaissetia nigra*, a scale insect species in the family Coccidae (NCBI accession: KY927598.1, KY924795.1). We estimated a phylogeny from the concatenated alignment (SYM+I+G4 substitution model) with both sets of markers. Additionally, we also estimated phylogenies for each marker alone (**Supplementary Figure 1**). Although the topology of phylogenies produced with each marker separately differed in some respects from each other, the number of losses of heterochromatic bodies was consistent across phylogenies.

### Microsatellite inheritance assay

To further explore whether paternal chromosomes are transmitted through sperm, we used microsatellite loci to assess transmission in two species of *Cystococcus* that differed in the presence of heterochromatic bodies in somatic cells of males (*Cystococcus echiniformis-*absent and *Cystococcus campanidorsalis-*present). *Cystococcus* females form large galls and male offspring develop within their mother’s gall. Once males mature, the female gives birth to daughters (first instar offspring) that disperse from the gall on the abdomens of their adult brothers (Semple *et al*., 2015). Therefore, it is possible to collect families from the field consisting of the gall, the female within her gall, and her offspring, and male offspring are easy to distinguish from female offspring because they are morphologically distinct at all developmental stages (Gullan and Cockburn 1986; Semple *et al*., 2015).

We determined gene transmission from mothers to sons in seven families of *C. campanidorsalis* and 13 families of *C. echiniformis* (**Supplementary Table 1**). Although we could not directly genotype the father/s, we were able to estimate if the genotypic contribution of the father/s was haploid (as expected under PGE) or diploid (as under Mendelian inheritance. We designed nine primer pairs for each species for polymorphic microsatellite loci from whole genome sequence data, using QDD software (Meglécz *et al*. 2014) to generate primers (**Supplementary Table 5**; see **Supplementary Methods** for primer design information). However, one primer for *C. campanidorsalis* gave inconsistent results so we did not analyse data from that locus, giving a total of eight loci for that species. We extracted 20μl DNA using the PrepGEM insect extraction kit (ZyGEM) from up to 20 male offspring from each family following manufacturer instructions, and a small portion of body wall tissue from the mother, and performed PCR reactions with the Type-it microsatellite PCR kit (Qiagen) (see **Supplementary Methods** for additional information on PCR procedure).

We analysed the microsatellite profiles of each individual in Geneious Prime (2020.2.3) with the microsatellite plugin, manually calling microsatellite peaks. We determined for the male offspring of each family how many alleles they had inherited from their mother and the inferred number of alleles they had inherited from their father (by examining the genotypes of all male offspring and excluding the mother’s genotype). Using the package lme4 in Rstudio (v.3.6.3) (Bates *et al*. 2014, R-Core-Team, 2020), we analysed whether males inherited a different number of alleles from their mother and father and whether the two species show different inheritance patterns, using a generalised linear mixed effects model with a binomial distribution. We were interested in the number of loci that exhibited haploid vs. diploid inheritance patterns in male offspring for each family. Therefore, we counted the number of microsatellite loci for which each family inherited one allele vs. two alleles from each parent, and combined the count into a vector for the response variable. We included the species and the source of alleles (maternal/paternal) as fixed effects, and an observation level random effect.

As the samples we analysed were field-collected, we did not have any information about whether females had mated once or multiply. Therefore, in order to determine whether cases where families had inherited more than one paternal allele were caused by the father being diploid and transmitting either allele to his offspring (i.e. no PGE), or by multiple males with PGE and with different genotypes mating with the same female, we examined the allele inheritance patterns for families that had inherited two paternal alleles for two or more microsatellite loci. We would expect that as long as the microsatellite loci were not located in close proximity to each other on the chromosome (note that *Cystococcus* species have at least 2n=18 chromosomes) that the allele inheritance profile for different microsatellite loci would be unrelated to each other in the case that the species does not exhibit PGE. However, in the case that the species does exhibit PGE and the female had mated with multiple males, we would expect the inheritance profile for different microsatellite loci to be related as offspring with the same father would inherit one set of alleles, and the offspring with a different father would inherit a different set of alleles. We conducted a chi-square test in Rstudio, with the number of individuals with each genotype as the observed frequencies, and the expectation that allele inheritance is random to determine if the offspring genotypes deviated from what would be expected if individuals in a family had one father that did not exhibit PGE.

### Expression of paternally derived alleles in males

Male mealybugs with heterochromatic bodies exhibit suppressed expression of paternally inherited chromosomes in somatic tissue (de la Filia *et al*., 2021). To determine whether somatic cells in males of *Cystococcus* without heterochromatic bodies express genes inherited from their father (i.e. exhibit diploid rather than haploid expression), we conducted an RNAseq analysis to examine allele expression in males and their mother and compare whether males exhibit uniparental (i.e. homozygous) expression of genes on a genome wide scale. We collected data from males of *C. campanidorsalis*, which have clear heterochromatic bodies and for which we expected expression from primarily maternally inherited genes, and males of *C. echiniformis*, which lack heterochromatic bodies and we therefore expected to show diploid (biparental) expression (see **Figure 2** for images of somatic cells). We collected galls for *C. echiniformis* and *C. campanidorsalis* from Queensland Australia in 2017, and preserved a small portion of somatic tissue from the female in each gall, and the whole body of two of her sons in a small volume of RNAlater for extraction. We were able to collect one family for *C. campanidorsalis* and three families for *C. echiniformis* (**Supplementary Table 1)**.

**Figure 2.**
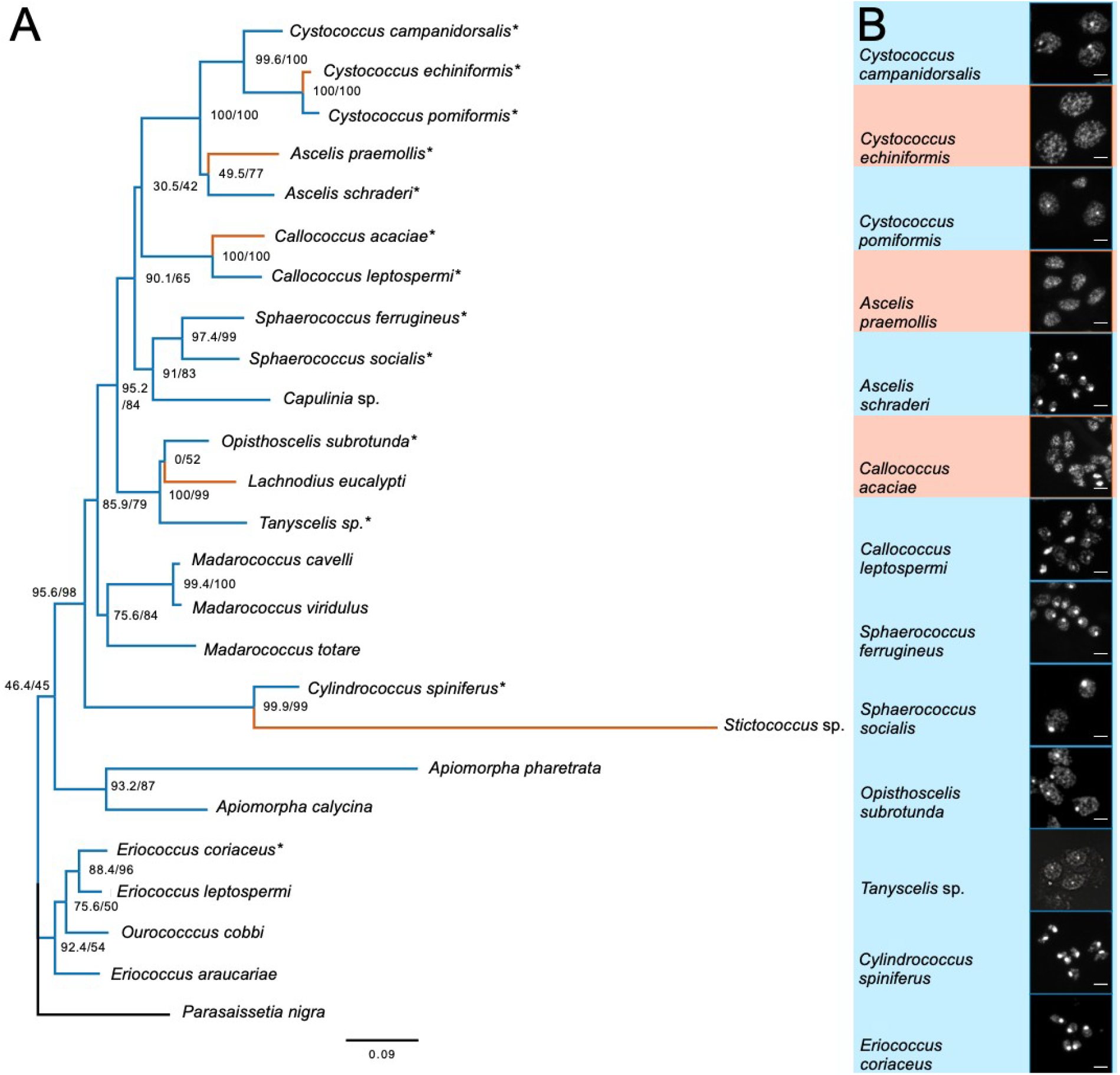
(**A**) Maximum likelihood phylogeny of eriococcid species for which we have data on whether heterochromatic bodies are present in males. Branches for species without heterochromatic bodies are coloured orange, while those with heterochromatic bodies are coloured blue. The phylogeny was generated with 18S and CO1 markers and an SYM+I+G4 substitution model with *Parasaissetia nigra* (Coccidae) included as the outgroup species (branch coloured black). Values at nodes show the SH-aLRT/ ultrafast bootstrap support with 1000 replicates. Species examined in this study are indicated with asterisks, whereas the occurrence of heterochromatic bodies for other species was taken from Brown (1967) or Brown (1977). (**B)** Images of somatic cells of males from eriococcid species examined in this study showing species with (blue) or without (orange) heterochromatic bodies.

We extracted RNA from the body wall of females using a Trizol:chloroform extraction and digesting residual DNA after extraction using DNAse I. Because the male samples were much smaller than the female tissue, we used a different RNA extraction procedure to account for the low yield from these samples. For males, we extracted RNA from the whole body using a modified Trizol RNA extraction procedure with the Purelink RNA Purification Kit (Promega), followed by amplification of the product with the Ovation RNAseq System V2 (Tecan) (See **Supplementary Methods** for full RNA extraction protocols). The samples were sequenced by Edinburgh Genomics, generating 150bp paired-end reads with TruSeq stranded mRNA-seq (or DNA) libraries, as appropriate.

We trimmed the sequence with fastp with settings --cut_by_quality5 --cut_by_quality 3 -- cut_window_size 4 --cut_mean_quality 20 (v 0.12.3) (Chen *et al*. 2018), then assembled a de novo transcriptome for both species with all libraries for that species using Trinity with parameters --full_cleanup and --SS_lib_type RF (v2.8.4) (Grabherr *et al*. 2011). We then conducted a BUSCO analysis (v3.0.2) (Waterhouse *et al*. 2018) on each species using the insecta_odb9 database (**Supplementary Table 6**). As de novo transcriptome assemblies can result in an unexpectedly high number of predicted transcripts, we used previously established approaches to filter the number of transcripts based on expression (Moghadam *et al*. 2013). We mapped the trimmed reads of each species to the appropriate transcriptome using RSEM with parameters --paired-end --bowtie2 (v1.3.1) (Li and Dewey 2011; Langmead and Salzberg 2012), then filtered out any transcripts which had an FPKM < 0.5. We then used the Trinity script get_longest_isoform_seq_per_trinity_gene.pl to retain the longest isoform for each transcript. We ran the TransDecoder pipeline included with Trinity, to retain only transcripts that had known homology in each transcriptome. We first used TransDecoder.LongOrfs with default parameters. We then conducted blastp searches of the TransDecoder.LongOrfs output against the SWISS-PROT database with an e-value cutoff of 1e-5, and hmmscan searches against the Pfam database (31.0) with default parameters. We predicted coding regions with transdecoder predict with parameters --single_best_only --retain_pfam_hits --retain_blastp_hits.

We mapped the trimmed reads to the resulting transcripts with Bowtie2 (v2.3.5.1) (Langmead and Salzberg, 2012), marked PCR duplicates with samtools markdup (v1.9) (Li *et* al. 2009), and called variants on the transcripts with freebayes (v1.3.1) (Garrison and Marth 2012), using all libraries for each species to call variants with parameters -w --standard-filters -C 5 --min-coverage 10. We used vcffilter from vcflib (v1.0.0_rc2) (Garrison, 2012) and retained SNPs with a depth of 10, a quality of 20, at least 2 alternate alleles on both strands, and an alternate allele balance from 0.2 to 0.8 in individuals called as heterozygous. We also filtered out multiallelic SNPs(1.1% and 2.0% of all SNPs in *C. campanidorsalis* and *C. echiniformis* respectively). We then used ASEReadCounter in GATK (v4.1.9.0) (McKenna *et al*. 2010) to count the number of reads supporting reference and alternate alleles at each SNP for each sample (i.e. for the reads from each individual separately) with the parameters --min-depth-of-non-filtered-base 40 --min-base-quality 20 --min-mapping-quality 40. We used Rstudio to determine the proportion of homozygous and heterozygous SNPs for each sample, filtering out SNPs in which the proportion of other bases (bases which were not the reference or alternate allele) was larger than 0.05. The vast majority of SNPs were retained after this filtering. We scored any SNPs with allele bias (proportion of reference to total allele count) between 0.2 and 0.8 as heterozygous, and any SNP with an allele bias less than 0.1 or greater than 0.9 as homozygous. We then compared the frequencies of homozygous and heterozygous SNPs between males and females for each species, with the expectation that if a significant part of the paternally-derived genome in males is silenced, then males should show an excess of homozygous expression compared to females (similar to de la Filia et al., 2021).

## Results

### Several eriococcid species have lost male somatic heterochromatization

Of the 13 species we examined, nine clearly exhibited heterochromatic bodies within somatic tissue of males (**Figure 2**). The appearance and size of heterochromatic bodies, however, varied between species and for different tissue within the same species: in some species heterochromatic bodies were clearly visible in some tissues but not visible in others, and some species had a higher proportion of nuclei that contained heterochromatic bodies compared to other species (**Figure 2; Supplementary Figure 2**). In *C. echiniformis, A. praemollis*, and *Callococcus acaciae*, we were unable to identify heterochromatic bodies in any individuals. Species which have lost male somatic heterochromatization, including *L. eucalypti* and *Stictococcus* sp. (Brown, 1977), are distributed across the Eriococcidae phylogeny (**Figure 2**). This suggests that the loss of somatic paternal chromosome heterochromatinization has occurred multiple times independently across the family.

### Male meiosis varies between species in the Ascelis/Cystococcus clade

We were able to examine spermatogenesis in six species and found that male meiosis varies substantially between Eriococcidae species. We identified pycnotic nuclei in “*Sphaerococcus” ferrugineus* and *Eriococcus coriaceus*, indicating that these species exhibit PGE (as noted by Brown (1967) for *E. coriaceus*) (**Supplementary Figure 3**). In *Ascelis praemollis*, which does not have heterochromatic bodies in the soma, sperm bundles occasionally had pycnotic nuclei associated with them, suggesting that this species exhibits PGE despite not having heterochromatic bodies (**Supplementary Figure 3**). However, we could not identify pycnotic nuclei in sperm bundles of any *Cystococcus* species we examined (**Figure 3; Supplementary Figure 3**), suggesting males might exhibit PGE with only one division in meiosis or might not exhibit PGE.

**Figure 3.**
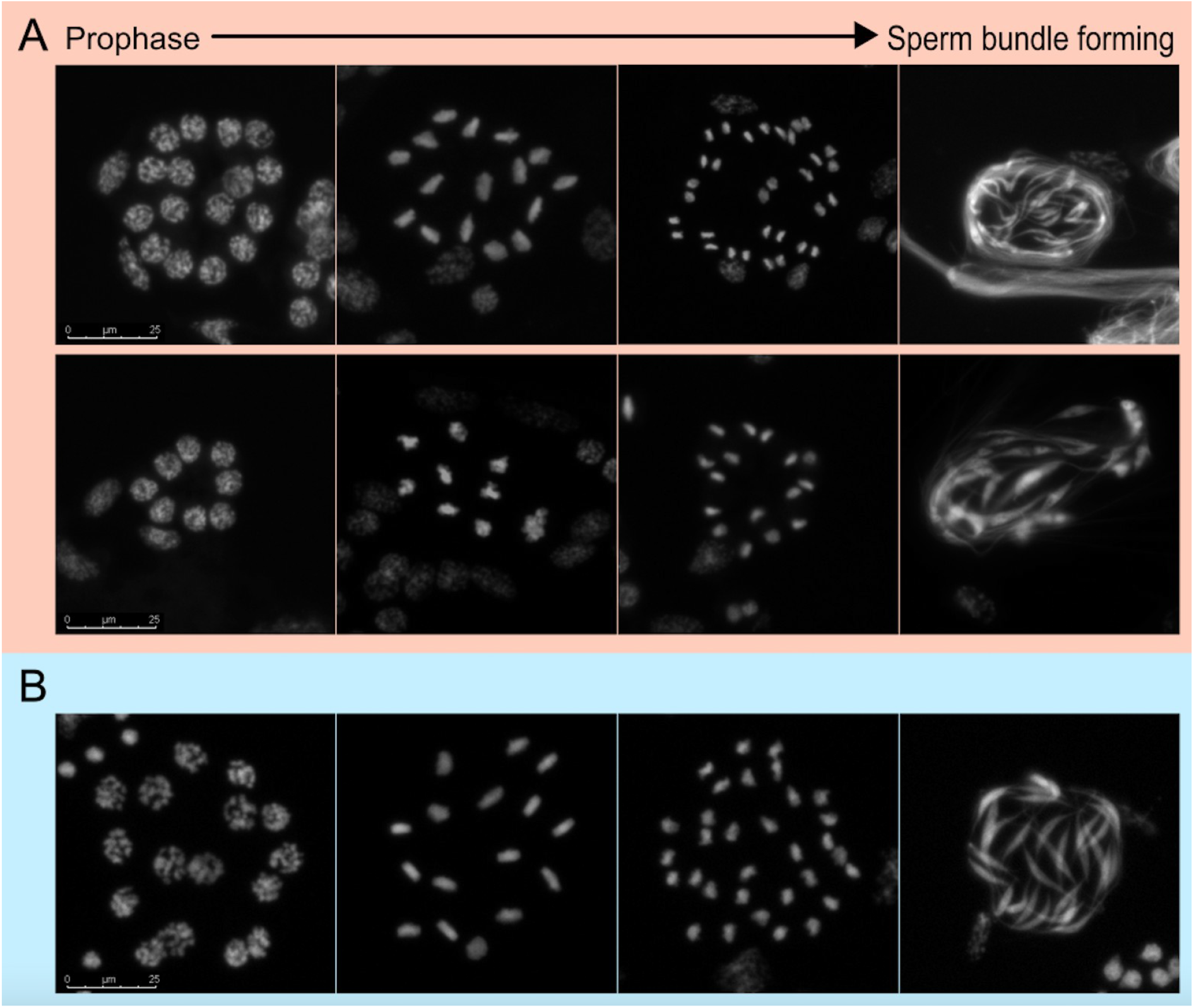
Meiosis in *Cystococcus echiniformis* (**A**), a species without heterochromatic bodies in somatic cells of males, and *C. campanidoralis* (**B)**, a close relative with heterochromatic bodies. Images show a sperm cyst containing nuclei throughout different stages of meiosis or following meiosis when sperm is forming. In both species pycnotic nuclei are absent following meiosis (last image in each panel; **Supplementary Figure 3**), and there is one division in meiosis, suggesting that both species exhibit PGE. However, there is likely variation in the number of nuclei per sperm cyst within *C. echiniformis* individuals, such that some sperm cysts start with eight nuclei and some sperm cysts start with 16 nuclei (first vs. second panel of **A**).

In all *Cystococcus* and *Ascelis* species examined in detail, we made sure to stain a variety of male developmental stages, such that we were confident that we were capturing the full meiotic process. Spermatogenesis in scale insects occurs within a cyst where several spermatogonia undergo meiosis concurrently. In *C. campanidorsalis*, we always observed 16 nuclei per sperm cyst at prophase and 32 sperm elongating after meiosis, indicating that *C. campanidorsalis* has one division in meiosis and Comstockiella PGE, with the elimination of all paternal chromosomes *prior* to meiosis. In *C. echiniformis*, sperm cysts most often had 16 nuclei in prophase sperm cysts, but in the same individuals we also observed a minority of sperm cysts with 8 nuclei at prophase. We most often observed 32 sperm following meiosis, but there were a few cases in which we observed 16 sperm following meiosis (**Figure 3; Supplementary Figure 4**). This could indicate that, in *C. echiniformis*, the number of nuclei in sperm cysts can vary between 8 and 16. In this case, *C. echiniformis* also exhibits PGE since there is only one division in meiosis, indicating that paternally inherited chromosomes are eliminated prior to meiosis. Alternatively, *C. echiniformis* may not exhibit PGE and may undergo two divisions in meiosis. We explore these possibilities further in the microsatellite analysis. In *C. pomiformis*, we were only able to view meiosis in prophase and after sperm bundles formed, but we found that this species also has some sperm cysts with 8 nuclei and some with 16 nuclei, and also has some sperm bundles with 16 sperm, and some with 32 (**Supplementary Figure 2**). Therefore, this species has a similar type of PGE to *C. echiniformis*. Finally, both *Ascelis* species have 8 nuclei in primary spermatids and 16 sperm forming in each sperm cyst after meiosis, indicating that both of these species have Comstockiella PGE and one division in meiosis (as noted for *A. schraderi* in Brown 1967) (**Supplementary Figure 5**).

### *Cystococcus* species exhibit allele inheritance patterns consistent with paternal genome elimination

To further investigate whether *Cystococcus* species exhibit PGE inheritance, we examined allelic inheritance patterns for 9 and 8 microsatellite loci for male siblings and their mother in *C. echiniformis* and *C. campanidorsalis* respectively. We analysed the genotype of broods of siblings with the expectation that if the species exhibits PGE, all the microsatellite loci would show similar patterns of inheritance within the brood, with the brood inheriting either one or two alleles from its mother (depending on whether she was homozygous or heterozygous at that locus) and only one allele from the father since males only transmit the maternally inherited allele to their offspring under PGE. However, as the samples were field collected, we did not have information about whether females had mated multiply or once, and therefore inferred this information from the data. On average, *C. echiniformis* offspring receive 1.13 alleles from their father and 1.40 alleles from their mother, while *C. campanidorsalis* offspring receive 1.09 alleles from their father and 1.67 alleles from their mother for the loci examined (**Figure 4**). The number of cases in which individuals inherited two vs. one allele from their parent differed for alleles inherited from the mother vs. the father (glmer: est=2.38, s.e= 0.50, z= 4.73, p<0.001) but allele inheritance patterns did not differ between *C. campanidorsalis* and *C. echiniformis* (glmer: est=0.63, s.e.=0.48, z=1.30, p=0.194) (**Supplementary Table 5**). This suggests that both species exhibit the same type of inheritance, and that both species likely exhibit PGE, as the number of alleles inherited from the father were less than the number inherited from the mother.

**Figure 4.**
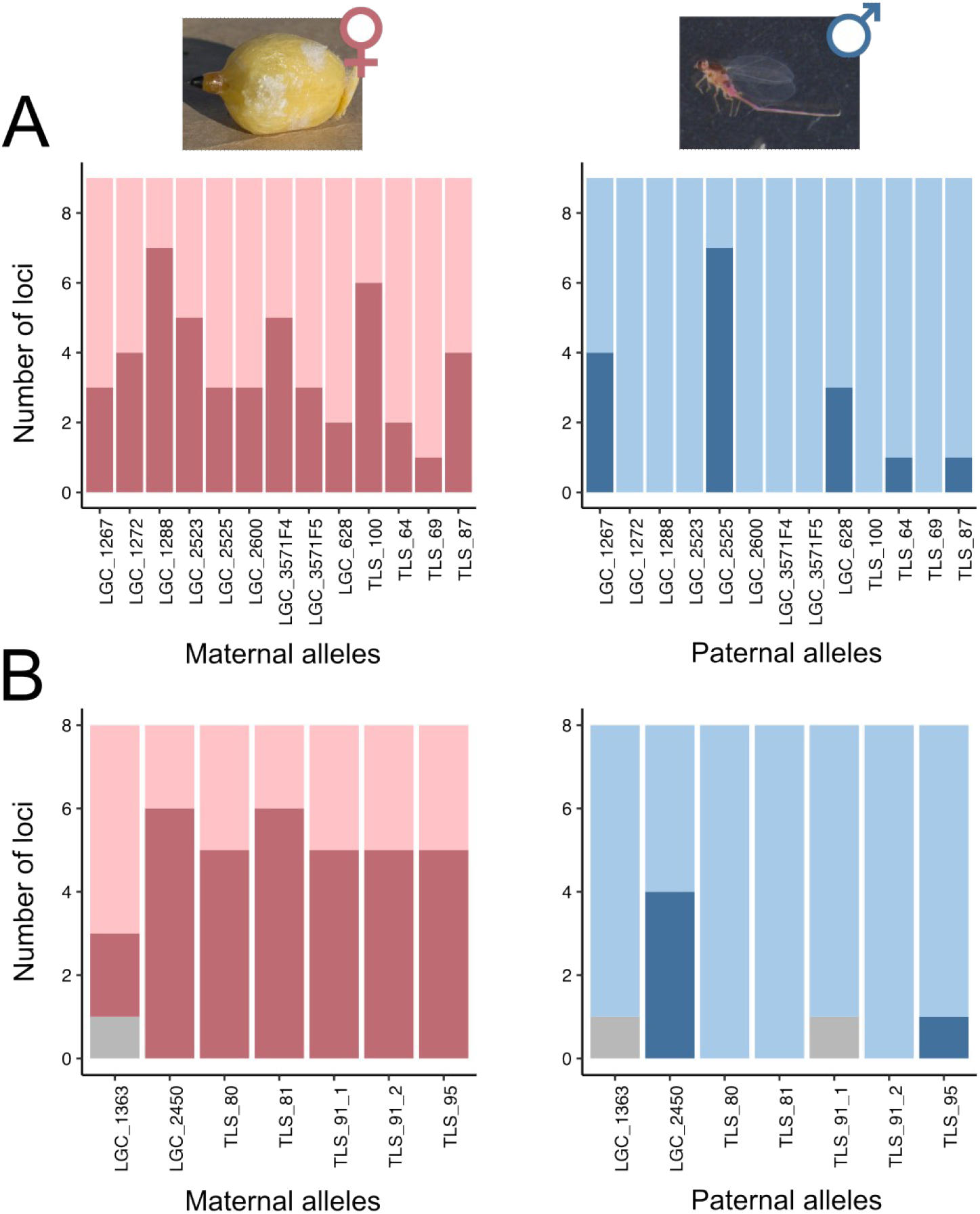
Inheritance patterns for male siblings from (**A**) *C. echiniformis* and (**B**) *C. campanidorsalis*. The inheritance patterns from the mother (pink) and the father (blue) are shown separately. The family ID vs. the number of microsatellite markers is shown in each plot, with the darker bar showing cases where families received two alleles from their parent and the lighter bar indicates cases where families received one allele (with grey bars indicating cases where the number alleles received was uncertain due to genotyping errors). For both species, siblings were more likely to inherit two loci from their mother than their father.

However, we did observe some instances where a family inherited two paternal alleles. For the cases where this involved more than one microsatellite locus, we investigated whether this was due to multiple mating (i.e., the mother mated with more than one male with PGE) or whether this was an indication of a lack of PGE transmission in that family. We were able to explore this in three families in *C. echiniformis* (families LGC_00628, LGC_01267, and LGC_02525), and one family in *C. campanidorsalis* (family LGC_02450). For all four of these cases, inheritance of alleles across microsatellite loci was not random (**Supplementary Table 6**, p<0.01 for all comparisons). Under PGE you would expect that males pass on the same haplotype to all their offspring (i.e. in PGE you expect linkage disequilibrium to be complete), so that if a female mated with two males (e.g., a 2-locus example with one male contributing AB and the other ab) her offspring would never inherit Ab or aB. However, a female mating with a single non-PGE male with an aAbB genotype would result in all possible allele combinations (**Supplementary Figure 6**). For the most part, we only observed two of the possible allele combinations in offspring, or in a few cases three (with the additional combination at low frequency so likely being the result of a genotyping error) (**Supplementary Figure 6)**. This indicates that in the cases we were able to examine, families that had inherited two paternal alleles more likely had mothers that mated multiple times than fathers that lacked PGE. Together these analyses support our cytogenetic results that both species have PGE transmission.

### Cystococcus males exhibit biparental gene expression

We also examined whether the loss of heterochromatic bodies from somatic cells of males indicates that males exhibit biparental gene expression, rather than predominantly maternal expression of genes, which we expect for males with heterochromatic bodies (de la Filia *et al*., 2021). To do so, we compared patterns of heterozygosity between females (mothers) and males (sons) of one *C. campanidorsalis* family and three *C. echiniformis* families, where we would expect an excess of homozygous SNPs in males if their paternal genome was either completely or partially silenced. We identified 39,439 biallelic SNPs in *C. campanidorsalis* and 30,184 biallelic SNPs in *C. echiniformis*. We filtered reads from each sample which did not meet quality filters, and excluded SNP positions from the final analysis which did not have sufficient depth of quality reads, resulting in slightly different numbers of SNPs that we considered for each individual (see **Supplementary Table 7**). *Cystococcus echiniformis* mothers exhibited heterozygous expression (20-80% expression of the non-reference allele) for 62.2%-63.5% of SNPs, while sons exhibited heterozygous expression for 59.0%-66.3% of SNPs (**Figure 5**; **Supplementary Table 7**). For *C. campanidorsalis*, the mother exhibited heterozygous expression for 77.0% of SNPs while the two sons exhibited heterozygous expression for 85.4% and 85.5% of SNPs. Overall, we found that sons exhibited biparental (heterozygous) allele expression patterns for a substantial number of SNPs in both *C. echiniformis* and *C. campanidorsalis*, suggesting that *C. campanidorsalis* males express alleles inherited from their father despite having heterochromatic bodies in somatic tissue.

**Figure 5.**
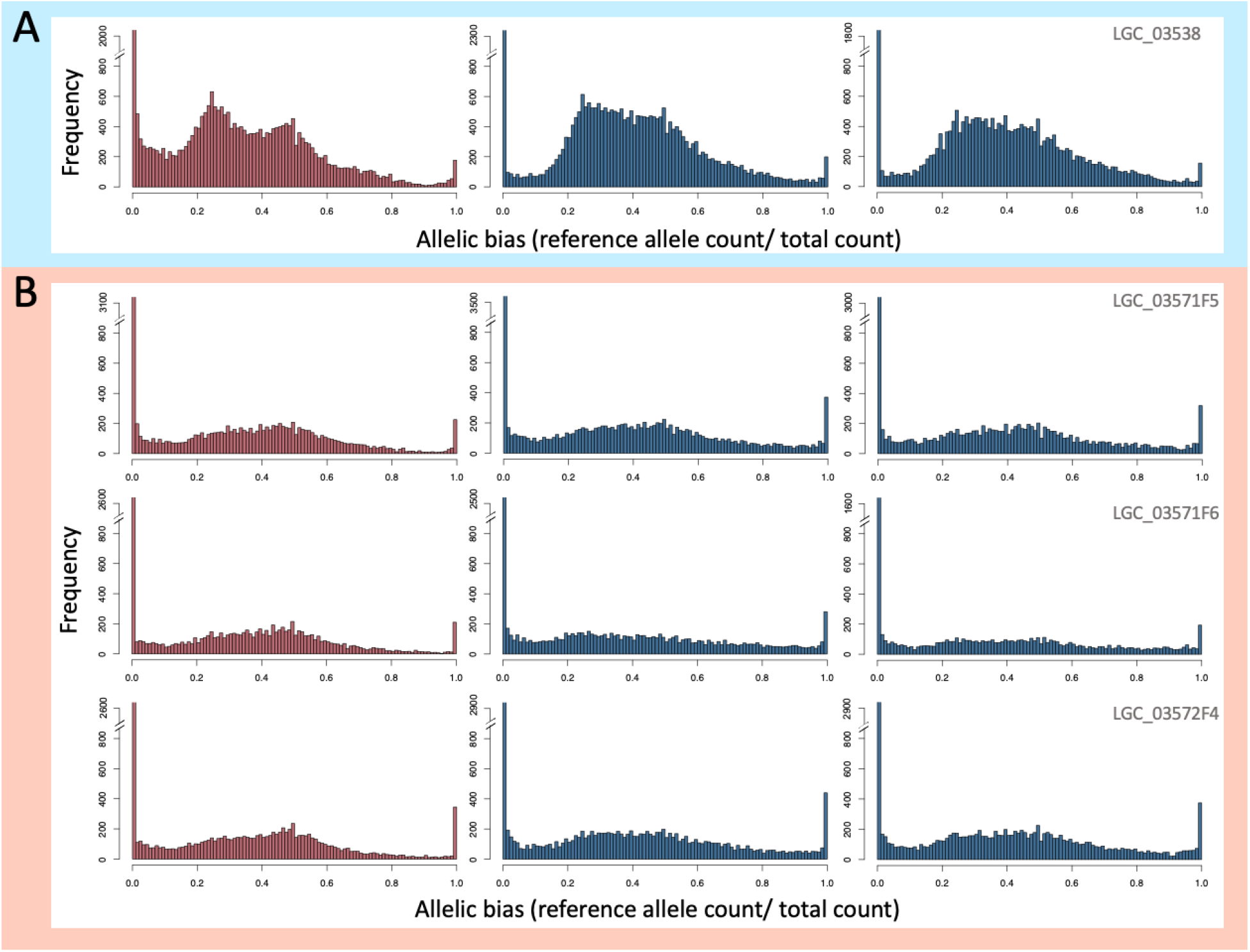
Histograms summarizing allele expression for mothers (red) and two sons (blue) for each family of *C. campanidorsalis* (**A**), and *C. echiniformis* (**B**). The frequency of the allelic bias (count of the called reference allele/ total allele count) for biallelic SNPs is shown, with alleles with homozygous (haploid) expression shown at either 0 or 1 and alleles with heterozygous expression shown from 0.2-0.8. In males with heterochromatic bodies, we would expect primarily homozygous (haploid) allele expression. Instead, mothers and sons have similar expression patterns in both species. The family ID is shown in the top right corner of each panel.

## Discussion

Asymmetric chromosome inheritance has evolved in numerous lineages (Ross et al., 2022). PGE and haplodiploidy are two common asymmetric systems, both involving exclusive transmission of maternally derived chromosomes in males. How such a system evolves, and whether reversions back to more typical Mendelian systems can occur, are currently unclear. For PGE in scale insects, conflict between the sexes over the transmission of the parental genomes through male offspring is thought to have led to its initial evolution, where the maternal genome displays meiotic drive (Brown 1964; Haig 1993). However, ongoing intragenomic conflict between the maternal and paternal genome within males is thought to have driven the transcriptional silencing of the paternal genome through heterochromatization, or the earlier elimination of paternal chromosomes (Herrick and Seger, 1999; Ross *et al*., 2010). Here, PGE with haploid males (e.g., Diaspidid PGE) would represent a scenario where the mother “won”, and monopolized transmission through males. However, in males that have retained chromosomes inherited from their father, there should be strong ongoing selection for paternally inherited chromosomes to regain transmission, which could have a number of consequences. First, it could drive the loss and/or silencing of the paternal genome through counteradaptations from the maternally inherited chromosomes. On the other hand, it may eventually lead to transmission of some or all of the paternally inherited chromosomes through sperm. However, given the significant mechanistic changes associated with the evolution of PGE (i.e., meiosis, spermatogenesis, ploidy, and sex determination), transitions back to Mendelian inheritance may be difficult or even impossible. We explored whether there is evidence for transmission of paternally inherited chromosomes through sperm (i.e., loss of PGE), or paternal chromosome expression in males using cytogenetic, genotyping and gene expression analyses in eriococcid scale insects: the only lineage in which PGE is thought to have been lost in several species.

Since eriococcid species ancestrally exhibit Comstockiella PGE, in which paternally inherited chromosomes are heterochromatized in males, one way of identifying changes in the presence/mechanism of PGE is by staining male somatic tissue to determine if heterochromatic bodies are present. We analysed samples from 13 species and found that, in addition to the two species that were previously found to have lost heterochromatic bodies (*Lachnodius eucalypti* and *Stictococcus* sp.), three other species within this clade have lost paternal somatic heterochromatization. Intriguingly, we also found evidence that the size of heterochromatic bodies, relative to the rest of the nuclei, differs both within and between species in this clade (**Figure 2**; **Supplementary Figure 2**). We currently do not understand why this variation may exist, but it may hint at ongoing conflict between maternal and paternal chromosomes in this family. As heterochromatic bodies contain all paternally inherited chromosomes, condensed into a heterochromatic ‘ball’, loss or variation in the size of the heterochromatic bodies might indicate that some or all paternally inherited chromosomes may be escaping transcriptional suppression in males and therefore may have a greater influence over transmission/ reproduction.

We found, through staining male reproductive tissue undergoing meiosis, that a lack of heterochromatic bodies does not necessarily mean a lack of PGE. Two species that lacked heterochromatic bodies, *C. echiniformis* and *A. praemollis*, have only one division in meiosis, suggesting that paternally inherited chromosomes are not passed on, but instead lost prior to meiosis (**Figure 3**; **Supplementary Figure 5**). Additionally, our microsatellite inheritance data showed *C. echiniformis* exhibits inheritance patterns consistent with PGE. Overall, this indicates that these species have transitioned to a different type of PGE, not previously reported in scale insects, but have not lost this type of reproduction entirely. This result opens debate over whether *L. eucalypti* and *Stictococcus* sp. really lack PGE, or whether earlier reports of meiosis in these species were misinterpreted. The evidence that *L. eucalypti* and *Stictococcus* sp. lacked PGE transmission was based on cytological data only (and with just a few available specimens) (Brown, 1977). It is therefore possible that these species exhibit a similar type of PGE to *C. echiniformis*, where variation in the number of cells per sperm cyst could be easily mistaken for evidence of two, rather than one division in meiosis. Further investigation of chromosome transmission in *L. eucalypti* and *Stictococcus* sp. is needed to corroborate earlier conclusions that PGE was lost. Unfortunately, this was not possible in our study: *L. eucalypti* is scarce, the only specimen we collected was an immature female so we were unable to study male meiosis and the *Stictococcus* sp studied previously was undescribed so it was not possible to follow up this work.

Herrick and Seger (1999) suggested that the elimination of paternal chromosomes just *prior* to meiosis evolved to prevent paternal chromosomes resisting maternal chromosome drive *during* meiosis. Therefore, variation in the characteristics of PGE may indicate ongoing conflict between the maternal and paternal halves of the genome in males.Our staining results indicate that in *C. echiniformis* and *A. praemollis*, the majority of paternal chromosomes are likely eliminated entirely before meiosis, as otherwise we would expect to see two divisions in meiosis rather than one (**Figure 3**; **Supplementary Figure 5, 7**). Although this may indicate a lack of opportunity for paternally inherited chromosomes to be transmitted to future generations, it is not conclusive that paternally transmitted chromosomes are never transmitted to offspring. In Comstockiella PGE systems, if most but not all of the paternal chromosomes are eliminated just prior to meiosis, there is one division in meiosis, with the remaining paternal chromosomes presumed to be eliminated afterwards (Nur, 1980), and in mealybugs, which have lecanoid PGE, occasional paternal chromosome leakage does occur through sperm (de la Filia *et al*., 2019). Therefore, there may be occasional leakage of paternally inherited chromosomes through sperm in eriococcid species. Although it would be fascinating to see if this ever happens, it would require a larger scale inheritance analysis than we undertook in this study, with larger sample sizes and more markers than we were able to use. It will also be challenging because unlike mealybugs which can be easily cultured under laboratory condition, the species considered in this study feed exclusively on trees of the family

## Myrtaceae, and are not straightforward to rear

We also found that males of both species exhibit substantial biparental gene expression. This result is surprising, as *C. campanidorsalis* has heterochromatic bodies, which are associated with substantial (although not complete) transcriptional suppression of paternally inherited chromosomes in males in the mealybug *Planococcus citri* (de la Filia *et al*., 2021). Variation in presence and size of heterochromatic bodies may be related to the expression levels of paternally derived chromosomes in males. The relative size of the heterochromatic bodies that we observed in all *Cystococcus* species were smaller than those observed in the mealybug *P. citri*, which could indicate that not all paternal chromosomes are part of the heterochromatic body. Alternately, the way that we assessed species for the presence of heterochromatic bodies, by scoring any species in which we observed heterochromatic bodies as possessing this trait, makes sense when scoring for the presence of PGE, but perhaps not when thinking about expression of paternally inherited chromosomes. For instance, in *C. pomiformis*, a close relative to *C. campanidorsalis*, we noted that some cells have heterochromatic bodies and some cells do not (**Supplementary Figure 2**). Therefore, perhaps not all tissues exhibit heterochromatic bodies in *C. campanidorsalis* resulting in a significant amount of biparental expression at the whole body level. Tissue specific heterochromatization has been previously noted in other scale insect species, including *P. citri*, where some somatic and germline tissues do not possess heterochromatic bodies (although it is not clear how this relates to expression in those tissues) (Nur, 1967; de la Filia *et al*., 2021).

The fact that males in *C. echiniformis* and *C. campanidorsalis* both express paternal chromosomes in somatic cells is intriguing. Eriococcid species evolved from an ancestor with somatic heterochromatization (Nur, 1980; Ross *et al*., 2010), suggesting that the ancestor likely showed limited expression of paternal alleles, like mealybugs (de la Filia *et al*., 2021). Therefore, re-evolving biparental expression from a uniparental chromosome expression system seems possible in PGE species, and leads to a transition from haploid to diploid male expression. Although the mechanism of sex determination in scale insects is unclear (as no species in this clade have heteromorphic sex chromosomes), somatic heterochromatization was also suggested to be an important aspect of the sex determination system (Buglia *et al*. 2009). Our results indicate that this may not be the case. However, our study was designed to assess broad patterns of expression, specifically whether males exhibited haploid or diploid chromosome expression on a whole genome scale. Because of this, there are still many questions to be answered about chromosome expression in *Cystococcus* species. For instance, are there any chromosomes/genes in which expression is strictly maternal and do all tissues exhibit the same expression patterns? How has the shift to biparental chromosome expression affected dosage compensation (i.e., overall gene expression levels), and do any related species have strictly maternal chromosome expression? Perhaps a good species to explore this last question in is *Ascelis schraderi*, which is closely related to *Cystococcus* species (Semple *et al*., 2015) and exhibits large heterochromatic bodies in most tissues (**Figure 2**).

The type of PGE found in *C. echiniformis*, with no heterochromatic bodies in somatic cells, was previously only known from head and body lice (de la Filia *et al*. 2018). Our results suggest that the evolution of PGE may have followed a similar trajectory in lice and scale insects. PGE has been characterised in few lice species (only head/body lice and *Liposcelis* booklice) (Hodson *et al*. 2017; de la Filia *et al*. 2018). However, all parasitic lice exhibit a peculiar type of meiosis in males (reviewed in White 1973) and, as head and body lice are nested within the monophyletic clade of parasitic lice to which booklice form the outgroup (Yoshizawa and Johnson 2010), the entire clade likely exhibits PGE. Booklice have a similar type of PGE to Lecanoid scale insects, with heterochromatic bodies in somatic cells of males (Hodson *et al*., 2017). The evolutionary trajectory seems to be similar in lice and scale insects, with heterochromatic bodies and PGE being found in basally branching clades and the more recent evolution of a loss of heterochromatic bodies and biparental gene expression in males, with PGE transmission dynamics. More in depth examination of the mechanism of PGE in both clades can provide insight into general trends in the evolution of PGE and asymmetric inheritance.

### Concluding remarks

Our results indicate that the mechanism of PGE is variable in scale insects, but there is little evidence for transitions from PGE to Mendelian inheritance. Also, the two scale insect species in which PGE was previously thought to be lost should be re-examined. Understanding more about the mechanism of reproduction in species with PGE helps us determine how likely it would be for PGE to transition back to Mendelian inheritance. For instance, in scale insects, we know that the transition to PGE must have involved a shift in how sex determination occurs in this lineage, as the ancestral XO sex determination system would not result in offspring of both sexes with PGE transmission dynamics. However, we still do not understand how sex determination occurs in scale insects and, therefore, whether it would be difficult to lose PGE transmission dynamics and still retain viable individuals that are able to reproduce in this lineage. This is also true in other lineages with PGE. Therefore, surveys of the characteristics of PGE and reproduction in taxonomically diverse species with asymmetric inheritance provide a valuable foundation from which to answer questions about the evolution of non-Mendelian inheritance.

Although we did not find evidence for a loss of PGE in eriococcid scale insects, we did find evidence for a previously unknown mechanism of PGE within scale insects. In *C. echiniformis*, males exhibited PGE transmission dynamics despite a lack of heterochromatic bodies, and in both *C. echiniformis* and *C. campanidorsalis*, we find that paternally inherited chromosomes are expressed in males, which was not previously thought to occur. Future work into whether the shift from haploid to diploid chromosome expression in *Cystococcus* species is complete, and how expression patterns compare to species with haploid expression, is needed. Eriococcidae are therefore an ideal clade of insects for studying how variability in epigenetic chromosome modification affects expression, and changes in ploidy and associated dosage compensation mechanisms evolve.

## Data availability statement

Scripts associated with this study are deposited on github (XX) and zenodo (XX). Sequences used in the phylogenetic analysis are deposited on NCBI (accession: XX) and RNAseq data is deposited under accession: XX.

## Acknowledgements

We would like to thank Andrés de la Filia and the Ross lab for comments on this manuscript and analysis advice. We would also like to thank the Cook lab for their hospitality and support.

