## Supplementary information for "Are asymmetric inheritance systems an evolutionary trap? Transitions in the mechanism of genome loss in the scale insect family Eriococcidae"

### Supplementary Methods

#### *Staining details*

The staining procedure was as follows: first, we dissected two individuals on a coverslip in freshly prepared cold 45% glacial acetic acid by spreading the body tissue into a thin layer with pins. We squashed the tissue firmly with a microscope slide, then briefly froze the slide in liquid nitrogen and removed the coverslip with a razor blade. We rehydrated the slide for 5 minutes in approximately 20µl 2X saline sodium citrate (SSC), then removed excess SSC with a kimwipe. We added approximately 20µl of Vectashield with DAPI (Vector Laboratories) over the tissue when the slide was nearly dry, added a coverslip and sealed the coverslip with nail polish. We stored slides in darkness at 4°C. We viewed slides on a Leica DM 2000 LED microscope with a DAPI filter and took images with Leica Application Suite software.

#### *Microsatellite primer design*

We extracted DNA from a portion of the body wall of an adult female using a CTAB DNA extraction protocol. For the DNA extraction, we first added 600µl of CTAB (Sigma-Aldrich) and 5µl of proteinase K, and incubated the sample overnight at 56°C. We then centrifuged the sample at 13,000rpm for 3 minutes and transferred the supernatant to a new vial. We then added 500 µL of chloroform, inverted the tube 10 times, then centrifuged the sample for 3 min (at 13,000 rpm) and transferred the supernatant to a new tube. We repeated the chloroform step a second time, then added 200µl of a binding buffer (573.18g GuHCL, 500ml NH4Ac pH6, 500ml $H_2O$), along with 200µl of 100% EtOH, after which we vortexed the mixture and transferred it into a spin column. Then we followed the manufacturer's instructions from the Isolate II genomic DNA kit (Bioline) from the first ethanol wash step, finally adding 50µl of EB buffer to elute the DNA.

For *C. campanidorsalis,* TruSeq DNA library preparation and Illumina 150 base-pair paired-end sequencing (HiSeq2500) was completed at Macrogen Inc. (Republic of Korea) using one sixth of a lane. Potential SSR loci were identified using the QDD2 pipeline software package (Meglécz *et al.*, 2009) and BLAST to ensure that none were within known coding regions. Using Primer3 (Untergasser *et al.*, 2012) within QDD2, we designed primer pairs to amplify fragments between 90 and 400 base pairs in length, with a melting temperature between 57 °C and 63 °C and defaults for other parameters.

For *C. echiniformis*, we sequenced the sample at Edinburgh Genomics with MiSeq (250bp paired end seq with 350bp inserts) to generate low-coverage whole genome sequence data. We trimmed reads with fastp with parameters --cut_by_quality5 --cut_by_quality3 --cut_window_size 4 --cut_mean_quality 20 (v 0.12.3) (Chen *et al.*, 2018), and assembled the reads using default settings with CLC assembly cell (v5.0.0, Qiagen). We used this assembly to generate primers with QDD (Meglecz *et al.*, 2014). For primer generation, we used the default setting with the exception that we set the minimum PCR product size to 120bp. We also did not do the optional contamination check step (step4 in

Meglecz *et al.* 2014). Instead, we chose simple trinucleotide microsatellites as the target regions, generated primer sets that produced amplicon sizes ranging from 120-300bp, and blasted the contigs these primers were found on to the nr nucleotide database on NCBI, excluding any primers from contigs that blasted to non-Metazoan species. For both species, we chose 24 primer pairs and used the nine that produced the most consistent signals when multiplexed in groups of three primer pairs per PCR reaction.

*Microsatellite PCR and thermocycling conditions*

For PCR reactions, we used the Type-it microsatellite PCR kit (Qiagen), following the manufacturers guideline with a few variations due to the use of M13 fluorescent primers (Schuelke, 2000). We combined the nine microsatellite primer sets for each species into three PCR reactions each containing three primer sets. We conducted PCR reactions with the Type-IT Microsatellite Kit (Qiagen) in a total of 15µl, with 7.5 µl Type-it Mastermix, 0.375µl of the forward primer mix (with each primer at a concentration of 2µM), 1.5µl of the reverse primer mix (also at 2µM), 0.5µl of the M13 fluorescent primer (6FAM or VIC, at 5µM), 3.625µl, and 1.5µl of DNA. The thermocycling conditions for the microsatellite PCR was as follows (following guidelines from Schuelke, 2000): 94°C x 5 min, 30 x (94°C x 30sec, 56°C x 45sec, 72°C x 45sec), 8 x (94°C x 30sec, 53°C x 45sec, 72°C x 45sec), 72°C x 10min. We genotyped the product through Edinburgh Genomics using the ABI 3730 DNA Analyzer system (ThermoFisher Scientific) with LIZ 500 as the size standard. If the genotyping was unsuccessful, we re-ran the PCR with each primer set separately and sequenced the product, and if the PCR was still unsuccessful, we excluded that locus for that individual from analyses.

We aimed to genotype 20 males and the mother of each family for each microsatellite loci. However, due to differences in the number of sons collected for each family, and some DNA extractions or primer sets not working on some males, we analysed fewer than 20 males for some families (or for some primers for a family). For family TLS_091, we analysed 13 males, for family TLS_087 we analysed 15 males, for families LGC_01363 and LGC_02525 we analysed 17 males, for family TLS_095 we analysed 18 males, and for family TLS_100 we analysed 19 males.

*RNA extractions for gene expression analysis*

For RNA extractions from females, we extracted RNA from a small amount of body tissue from the females as we did not want contamination from germ tissue in the sample. We first took the female tissue out of RNAlater and rinsed it briefly in sterile 1X PBS before adding 50µl of Trizol to the sample and crushing the sample with a micropestle. We then added 950µl of Trizol and 200µl of chloroform, shook the sample by hand, transferred the supernatant to a new tube, added another 200µl of chloroform, and transferred the supernatant into a new tube for a second time. We then added 500ul of isopropanol and 1µl of linear acrylamide and stored the sample overnight at -20°C. The next day, we centrifuged the sample at 4°C for 15 min, removed the isopropanol from the samples, and added 1ml of freshly prepared 70% EtOH. We inverted the sample several times, centrifuged at 4°C for 15 min (13,000 rpm), and removed the EtOH with a pipette. We did the EtOH cleaning step twice for each sample. We dried the pellet at room temperature, then resuspended the pellet in 40ul distilled $H_2O$. We then performed a gDNA digestion, adding 1µl 10X reaction buffer with $MgCl_2$ (ThermoScientific), 1µl DNase I, 8µl water, and 0.25µl of RNase inhibitor for every 1µl of RNA in the sample. We heated the sample at 37°C for 30 min, then added 1µl 50uM EDTA and heated at 65°C for 10 minutes. Finally, we used the RNA Clean & Concentrator kit (Zymo Research) following the manufacturer instructions and eluting the samples in 60ul of $H_20$.

For the RNA extractions from male samples, we used the PureLink RNA Mini Kit (ThermoFisher Scientific), using a slightly modified protocol. We first briefly rinsed the samples in 1X PBS, then added 50µl Trizol into each sample and crushed the tissue with a micropestle. We then added 350µl of Trizol, briefly microcentrifuged the tubes, and transferred the supernatant to a clean tube. We added 80µl of BCP (1-Bromo-3-chloropropane), shook the sample by hand for 15 sec and incubated on ice for 3 min. We then centrifuged the samples at 4°C for 15 min (13,000rpm for all centrifugation steps). We transferred the supernatant to a new tube and added an equal volume of freshly prepared EtOH, mixing the tube by vortexing. We transferred the supernatant to a spin column and centrifuged the sample for 30 sec. We discarded the flow through and added 350µl of Wash Buffer I to the sample. We centrifuged for 30 sec, then added 80ul of the DNase mixture (made of 8µl 10X DNase I reaction buffer, 10µl resuspend DNase I, and 62µl of RNase-free water) onto the membrane of the spin column. We incubated this mixture for 15 min at room temperature, then added 350µl of Wash Buffer I to the sample and centrifuged for 30sec. We then placed the spin cartridge into a new collection tube, added 500µl of Wash Buffer II, and centrifuged the sample for 30sec. We repeated this step once, then centrifuged the sample for an extra minute to dry the membrane. We placed the spin cartridge into a clean eppendorf tube, added 30µl RNase-free water to the membrane, and incubated the sample for 1min. We centrifuge the sample for 2 min and stored the sample at -80°C. We performed a cDNA amplification of the male samples as the yield of RNA from these samples was small due to their small size. In order to do this, we used the Ovation RNAseq System V2 (Tecan), following the manufacturer protocol.

### Supplementary Tables/ Figures

**Supplementary Table 1**. Sample collection Information for samples used in this study.

| Species | Purpose | Collection date | Sample ID | Collection location | Latitude | Longitude | Collector |
|---|---|---|---|---|---|---|---|
| *Cystococcus campanidorsalis* | Microsatellite | 2013 | TLS_080F1 | | -27.45281 | 152.23029 | TLS |
| | Microsatellite | 2013 | TLS_081F2 | | -27.45281 | 152.23029 | TLS |
| | Microsatellite | 2013 | TLS_091F1 | | -24.39534 | 151.04507 | TLS |
| | Staining, microsatellite | 2013 | TLS_091F2 | | -24.39534 | 151.04507 | TLS |
| | Staining, microsatellite | 2013 | TLS_095F1 | | -24.4501 | 150.94395 | TLS |
| | Phylogeny | 26.iv.2008 | LGC_00847 | Crows Nest National Park, Qld | -27.2599 | 152.1172 | LGC |
| | Microsatellite | 14.xii.2009 | LGC_01363 | Giles Rd, Redland Bay, Qld | -27.3652 | 153.1656 | LGC |
| | Staining, microsatellite | 23.ii.2014 | LGC_02450F2 | Allies Creek State Forest, Qld | -25.95579 | 151.202625 | LGC |
| | RNAseq | 7.v.2017 | LGC_03538 | Lockyer Nat. Park, Qld | -27.4791 | 152.28159 | CNH |
| *Cystococcus echiniformis* | Microsatellite | 19.x.2013 | TLS_064 | | -27.72208 | 142.82253 | TLS |
| | Staining, microsatellite | 19.x.2013 | TLS_069 | 39km W of Thargomindah, QLD | -27.81193 | 143.51685 | TLS |
| | Microsatellite | 2013 | TLS_087F1 | | -24.58807 | 148.88364 | TLS |
| | Staining, microsatellite | 2013 | TLS_100 | | -26.07543 | 152.39366 | TLS |
| | Microsatellite | 12.ix.2006 | LGC_00628 | 12 km W of Herberton, Qld | -17.2248 | 145.1816 | LGC |
| | Phylogeny, microsatellite | 1.x.2009 | LGC_01267 | Town Lookout near Timber Creek, NT | -15.3846 | 130.2733 | LGC |
| | Microsatellite | 3.x.2009 | LGC_01272 | Keep River National Park, NT | -15.4501 | 129.0509 | LGC |
| | Microsatellite | 1.x.2009 | LGC_01288 | Joe Creek, Gregory National Park, NT | -15.3625 | 131.0442 | LGC |
| | Staining, microsatellite | 30.viii.2014 | LGC_02523F2 | Valley of Lagoons Rd, Qld | -18.515833 | 144.7836 | LGC |
| | Staining, microsatellite | 31.viii.2014 | LGC_02525 | Einasleight-Forsayth Rd, Qld | -18.54877 | 143.94232 | LGC |
| | Microsatellite | 20.ix.2014 | LGC_02600 | Homevale NP, Qld | -21.4078 | 148.5058 | LGC |

| | | | | | | | |
|---|---|---|---|---|---|---|---|
| | Microsatellite, RNAseq | 28.v.2017 | LGC_03571F4 | Hawkwood Rd, SW of Munduberra, Qld | -25.6956 | 150.9733 | CNH |
| | Microsatellite, RNAseq | 28.v.2017 | LGC_03571F5 | Hawkwood Rd, SW of Munduberra, Qld | -25.6956 | 150.9733 | CNH |
| | RNAseq | 29.v.2017 | LGC_03572F4 | Burnett Hwy, E of Gayndah, Qld | -25.6113 | 151.662 | CNH |
| *Cystococcus pomiformis* | Staining | 30.viii.2014 | LGC_02534 | Gregory Developmental Rd, Qld | -18.9961 | 144.695 | LGC |
| | Staining | 31.viii.2014 | LGC_02530 | Georgetown-Mt Garnet Rd, Qld | -18.8494 | 144.4247 | LGC |
| | Staining | 2.ix.2014 | LGC_02536 | Einasleigh-Forsayth Rd, Qld | -18.5447 | 143.8775 | LGC |
| | Phylogeny, Staining | 18.ix.2004 | LGC_00267 | Palm Grove Caravan Park, WA | | | LGC&MDC |
| *Ascelis praemollis* | Staining | 17.ix.2014 | LGC_02583 | Wild Rivers Caravan Park, Herberton | -17.22 | 145.2318 | LGC |
| | Staining | 14.x.2015 | LGC_02899 | 60 Bretons Rd, Crohamhurst, Qld | -26.802 | 152.87 | LGC |
| | Phylogeny | 16.vii.1994 | Asc1 | 391 Fisherman's Reach Rd, Stuarts Point | -30.49 | 153 | LGC |
| *Ascelis schraderi* | Staining | | PJM_00515 | School Road, Yeerongpilly, Qld | -27.521323 | 153.021189 | PJM |
| | Phylogeny | 26.xii.2006 | LGC_00718 | nr Trial Bay, South West Rocks, NSW | -30.52 | 153.03 | LGC |
| *Callococcus acaciae* | Staining | 4.x.2014 | LGC_02618 | Ku-ring-gai Chase NP, NSW | -33.673 | 151.134 | LGC |
| | Staining | 3.x.2014 | LGC_02614 | Red Hill reserve, Oxford Falls, NSW | -33.741 | 151.253 | LGC |
| | Phylogeny | 1.xi.2007 | LGC_01124 | O'Connor, ACT | | | MDC |
| *Callococcus leptospermi* | Staining | 4.xi.2014 | LGC_02626 | Myall Lakes NP, NSW | -32.5 | 152.285 | LGC |
| | Staining | 3.x.2014 | LGC_02612 | Elvina walking track carpark, NSW | -33.643 | 151.262 | LGC |
| | Phylogeny, Staining | 24.xii.2016 | LGC_03410 | St John's Wood Rd, Blairgowrie, Vic | | | LGC |
| *Cylindrococcus* sp. | Staining | 2017 | LGC_01956 | Burrabaranga Rd, Durikai State Forest, Qld | -28.26589 | 151.54376 | GPH |
| | Staining | | Fresh | Brisbane | | | LGC |

| | | | | | | | |
|---|---|---|---|---|---|---|---|
| | Phylogeny | 26.ix.2006 | LGC_00657 | Heathlands Resource Reserve, Qld | -11.38066 | 142.44216 | LGC |
| *Eriococcus coriaceus* | Staining | 1.vii.2014 | LGC_02489 | Bellangry State Forest, NSW | -31.288056 | 152.508611 | LGC |
| | Staining | v.2017 | Fresh | Colony on UQ, St. Lucia Campus, Qld | | | CNH |
| *Sphaerococcus ferrugineus* | Staining | 13.ix.2014 | LGC_02572 | Mt Emerald, near Tolga, Qld | -17.2103 | 145.4326 | MDC |
| | Phylogeny | 16.x.2006 | LGC_00685 | UQ, St Lucia Campus, Qld | -27.5005767 | 153.0152758 | LGC |
| *Sphaerococcus socialis* | Staining | 5.x.2010 | LGC_01654 | Chester Pass Rd, WA | -34.562166 | 118.00819 | MDC |
| *Opisthoscelis subrotunda* | Staining | 12.vi.2010 | LGC_01426 | Mt Tibrogargan, Qld | -26.55435 | 152.5652 | LGC |
| | Phylogeny | 8.ii.2004 | LGC_00099 | Midland Hwy, c. 27 km E of Shepparton, Vic | -36.26 | 145.42 | PJG |
| *Tanyscelis* sp. | Staining | 16.ix.2014 | LGC_02577 | Herberton-Irvinebank Rd, Qld | -17.38856 | 145.34669 | LGC |
| *Tanyscelis convexa* | Phylogeny | 17.x.2003 | LGC_00043 | Kennedy Hwy, 45 km SW of Mt Garnet, Qld | -16.5805 | 144.5144 | LGC&MDC |

**Supplementary Table 2.** Summary of samples stained and the results of cell staining for somatic cells as well as male tissue undergoing meiosis. For *Tanyscelis sp.* and *Capulinia jaboticabae,* we used sequence from a related species in the same genus in the phylogeny. We got the sequence for *Parasaissetia nigra* from NCBI (Accession: KY927598.1, KY924795.1). HB = heterochromatic body.

| Species | Reference | # | Life stage/ sex examined | Somatic HB | Meiosis stages examined | Number cells/ sperm cyst | Number sperm/ bundle | Pycnotic nuclei | Loci for phylogeny |
|---|---|---|---|---|---|---|---|---|---|
| *Cystococcus campanidorsalis* | this study | 20 | male nymphs/ pupae | Yes | all | 16 | 32 | No | 18S, COI |
| *Cystococcus pomiformis* | this study | 30 | male nymphs/pupae/ adults | Yes | Prophase/ sperm bundles | 8 or 16 | 16 or 32 | No | 18S, COI |
| *Cystococcus echiniformis* | this study | 25 | male nymph/ pupae/ adults | No | all | 8 or 16 | 16 or 32 | No | 18S, COI |
| *Ascelis praemollis* | this study | 6 | male nymphs/pupae | No | all | 8 | 16 | No/ few | 18S, CO1 |
| *Ascelis schraderi* | this study, Brown 1967 | 8 | male nymphs/pupae | Yes | Prophase/ metaphase | 8 | 16 | No | 18S, COI |
| *Callococcus acaciae* | this study | 11 | mixed sex crawlers/ early male nymphs | No | | | | | 18S, COI |
| *Callococcus leptospermi* | this study | 11 | mixed sex crawlers/ early male nymphs | Yes | | | | | 18S, COI |
| *Cylindrococcus sp.* | this study | 22 | mixed sex crawlers/ male nymphs and adults | Yes | | | | | 18S, COI |
| *Eriococcus coriaceus* | this study, Brown 1967 | 16 | mised sex crawlers/ male pupae | Yes | Sperm bundles | | | Yes | 18S, COI |
| *Sphaerococcus ferrugineus* | this study | 3 | male nymphs/ pupae | Yes | Sperm bundles | | | Yes | 18S, COI |
| *Sphaerococcus socialis* | this study | 11 | mixed sex crawlers/ male nymphs | Yes | | | | | 18S, COI |
| *Opisthoscelis subrotunda* | this study | 32 | mixed sex eggs/ crawlers/ male pupae | Yes | | | | | 18S, COI |
| *Tanyscelis* sp. | this study | | male nymphs | Yes | | | | | 18S, COI |
| *Apiomorpha pharetrata* | Brown, 1967 | | | Yes | | | | | 18S |
| *Apiomorpha calycina* | Brown, 1967 | | | Yes | | | | | 18S, COI |
| *Apiomorpha* (45 species) | Cook, 2000; 2001 | | | Yes | | | | | |

| | | | |
|---|---|---|---|
| *Capulinia jaboticabae* | Brown, 1967 | Yes | 18S, COI |
| *Capulinia orbiculata* | Brown, 1967 | Yes | |
| *Casuarinaloma leaii* | Brown, 1967 | Yes | |
| *Cylindrococcus spiniferus* | Brown, 1967 | Yes | |
| *Eriococcus abditus* | Brown, 1967 | Yes | |
| *Eriococcus araucariae* | Brown, 1967 | Yes | 18S, COI |
| *Eriococcus detectus* | Brown, 1967 | Yes | |
| *Eriococcus lecanioides* | Brown, 1967 | Yes | |
| *Eriococcus mimus* | Brown, 1967 | Yes | |
| *Eriococcus rata* | Brown, 1967 | Yes | |
| *Eriococcus rhodomyrti* | Brown, 1967 | Yes | |
| *Lachnodius eucalypti* | Brown, 1977 | No | 18S, COI |
| *Eriococcus leptospermi* | Brown, 1967 | Yes | 18S, COI |
| *Madarococcus cavelli* | Brown, 1967 | Yes | 18S |
| *Madarococcus totarae* | Brown, 1967 | Yes | 18S |
| *Madarococcus viridulus* | Brown, 1967 | Yes | 18S |
| *Ourococcus cobbi* | Brown, 1967 | Yes | 18S |
| *Phloeococcus loriceus* | Brown, 1967 | Yes | |
| *Tanyscelis convexa* | Brown, 1967 | only in germ tissue | |
| *Stictococcus* sp. | Brown, 1977 | No | 18S |
| *Parasaissetia nigra* (outgroup) | | Partheno-genetic | 18S,CO1 |

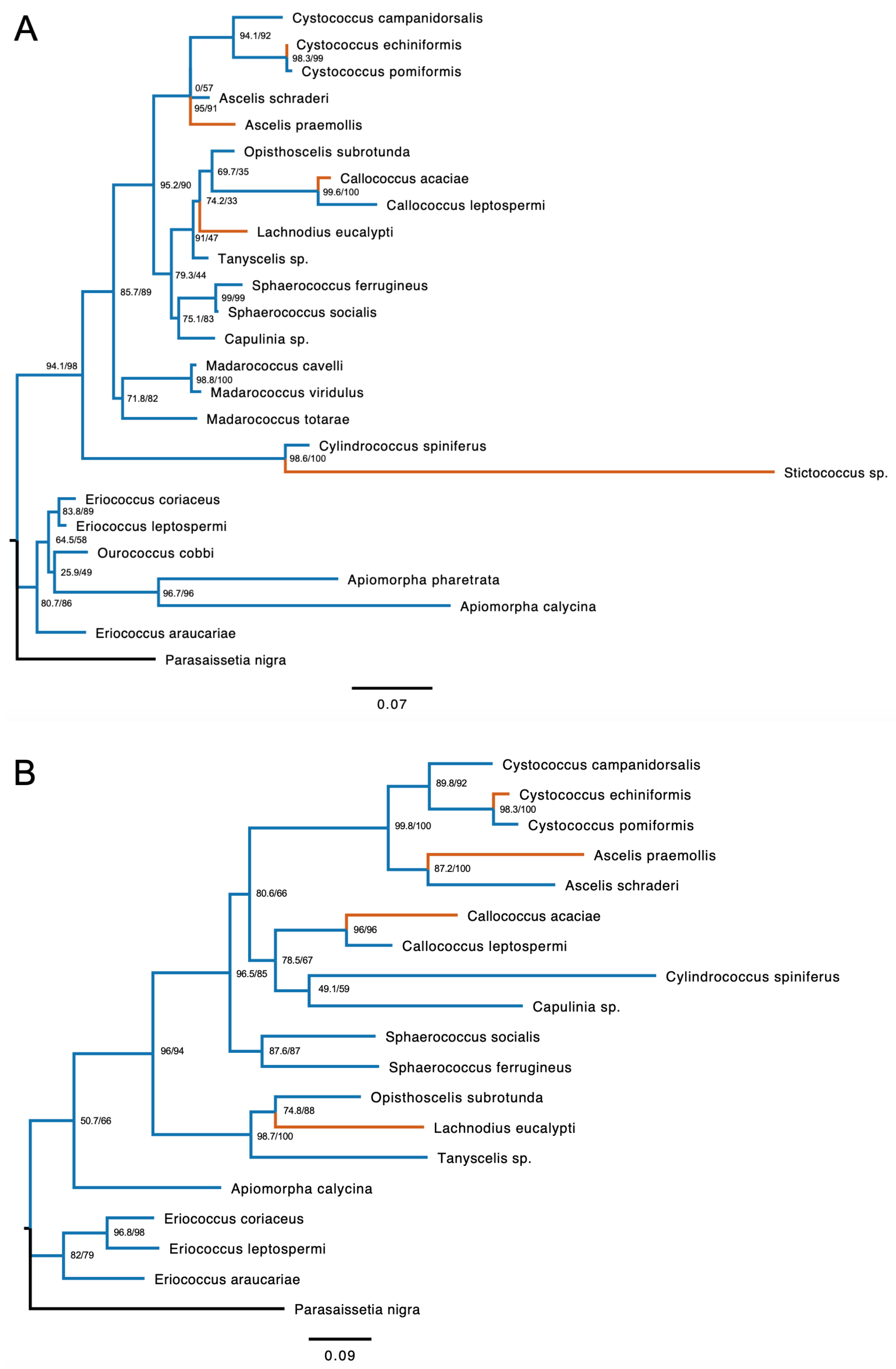


**Supplementary Figure 1**. Phylogeny of Eriococcidae species using an approximately 650 bp region of the nuclear gene 18S (**A**), and an approximately 600bp region of the mitochondrial gene CO1 (**B**). *Stictococcus* sp., *Apiomorpha pharetrata*, and *Ourococcus cobbi* have only 18S sequence available, and so are not included in (**B**).

**Supplementary Table 3**. Primer information for the microsatellite inheritance study, including primer sequences, the expected size of the PCR product for each primer pair, and which PCR panel each primer set belonged to. The M13 fluorescent primer was added into all PCR reaction mixes with a fluorescent tag to fluorescently label PCR products.

| Species | Primer name | Forward primer sequence | Reverse primer sequence | Expected size | PCR panel |
|---|---|---|---|---|---|
| *C. campanidorsalis* | cyst_msat_08 | GTAAAACGACGGCCAGAATTTCCATTCGGTAGTAGG | CAGTACAGTTACCATTCCACAG | 150 | 1 |
| | cyst_msat_09 | GTAAAACGACGGCCAGAAGGGAATTTAATGGTATGC | GAGGTTGGTGGTTCTAAGTG | 150 | 2 |
| | cyst_msat_12 | GTAAAACGACGGCCAGCGATGCGTTGAATATTAGTG | ATGTTCCGTGACCTAACTTG | 194 | 3 |
| | cyst_msat_14 | GTAAAACGACGGCCAGGAACCAAGCCAATAATGATC | GCAGCCCAAATATGAAGAG | 203 | 1 |
| | cyst_msat_16 | GTAAAACGACGGCCAGAAAGGTTGAAGGGTAGTGG | AACGGGAAATGTAAATTGAG | 217 | 1 |
| | cyst_msat_17 | GTAAAACGACGGCCAGAAATTATGGCCTTGAGTTG | TTAGGTGCCTCATACGTCAG | 221 | 3 |
| | cyst_msat_18 | GTAAAACGACGGCCAGTTGAGTGCGTAATTGAATTG | CACAGGCAGGTCTCTTAAAG | 222 | 2 |
| | cyst_msat_21 | GTAAAACGACGGCCAGGGAAGTGAATTTCGACGTAG | TTGCCCACCTACAGTAGTTC | 261 | 3 |
| *C. echiniformis* | c.ech_msat77 | GTAAAACGACGGCCAGCCCTATACCACGTTCGACCA | CCCTATACCACGTTCGACCA | 152 | 4 |
| | c.ech_msat79 | GTAAAACGACGGCCAGAAGCGTGTCTTCGTCTCGAT | AAGCGTGTCTTCGTCTCGAT | 124 | 5 |
| | c.ech_msat80 | GTAAAACGACGGCCAGTATTGCGCTATCTCATCGGA | TATTGCGCTATCTCATCGGA | 175 | 6 |
| | c.ech_msat89 | GTAAAACGACGGCCAGTTGACAAGAGCGAGAATTTCC | TTGACAAGAGCGAGAATTTCC | 238 | 4 |
| | c.ech_msat61 | GTAAAACGACGGCCAGGTCGTTAACCGATGGCAGAC | GTCGTTAACCGATGGCAGAC | 206 | 6 |
| | c.ech_msat97 | GTAAAACGACGGCCAGGGAATTCTATGCGAGGTTGC | GGAATTCTATGCGAGGTTGC | 234 | 6 |
| | c.ech_msat99 | GTAAAACGACGGCCAGCCACACCTCTTCGAAAGTCC | CCACACCTCTTCGAAAGTCC | 197 | 4 |
| | c.ech_msat102 | GTAAAACGACGGCCAGGGTCTTCCACGGATCAGTAGTT | GGTCTTCCACGGATCAGTAGTT | 141 | 5 |

| | | | | |
|---|---|---|---|---|
| c.ech_msat104 | GTAAAACGACGGCCAGGG GAGTCTTTACCACCTACGAT | GGGAGTCTTTA CCACCTACGAT | 233 | 5 |
| M13 Fluorescent tail | GTAAAACGACGGCCAG | N/A | | |

**Supplementary Table 4**: BUSCO results for de novo transcriptomes for *Cystococcus campanidorsalis* and *C. echiniformis*. Both species have more than 95% of the single copy orthologs expected to be present in insect genomes.

| | *Cystococcus campanidorsalis* | | *Cystococcus echiniformis* | |
|---|---|---|---|---|
| Complete | 1591 | 95.90% | 1589 | 95.80% |
| Single copy | 95 | 5.70% | 113 | 6.80% |
| Duplicated | 1496 | 90.20% | 1476 | 89.00% |
| Fragmented | 21 | 1.30% | 21 | 1.30% |
| Missing | 46 | 2.80% | 48 | 2.90% |
| Total | 1658 | | 1658 | |

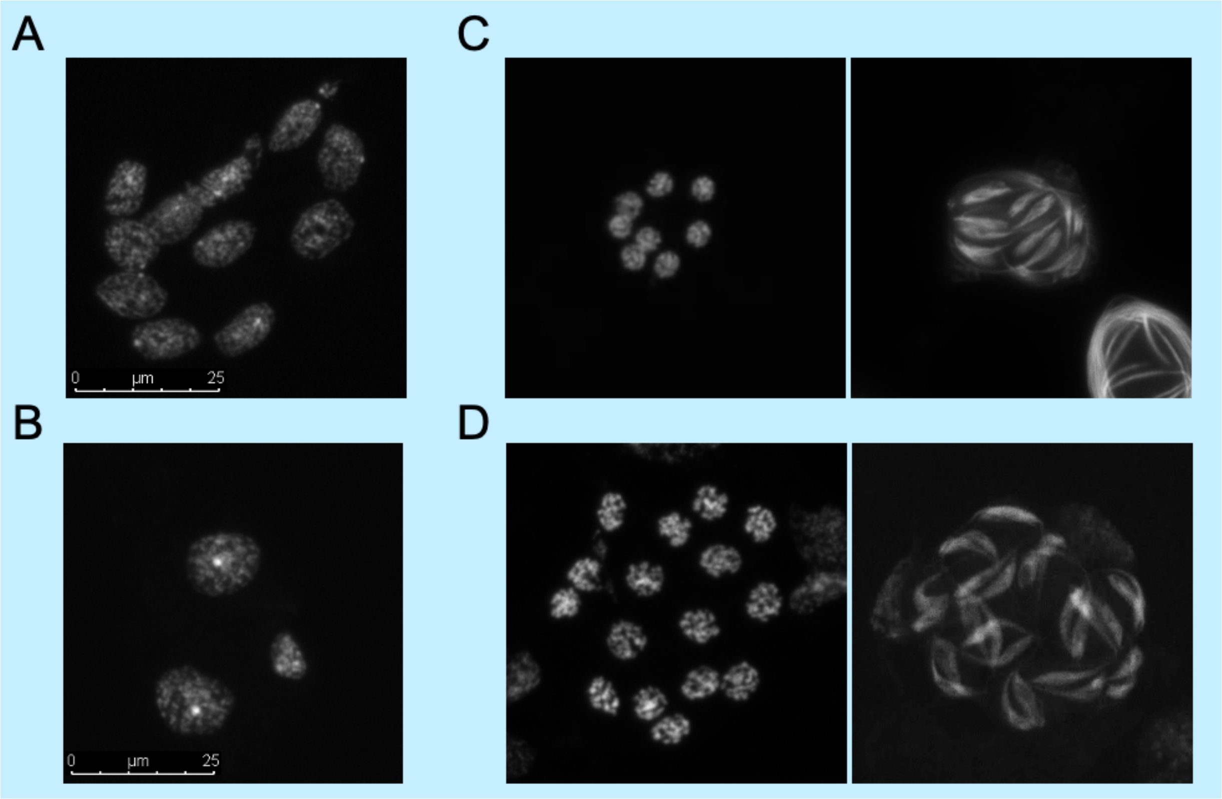


**Supplementary Figure 2**. Meiosis in *C. pomiformis*. (**A)** and **(B)** show somatic cells, (**A)** without (or with small) heterochromatic bodies and (**B)** with clear small heterochromatic bodies. (**C)** shows a sperm cyst with 8 nuclei and 16 sperm at the end of meiosis, while (**D)** shows a sperm cyst with 16 nuclei and 32 sperm at the end of meiosis. *Cystococcus pomiformis* likely exhibits Comstockiella PGE with only one division in meiosis, but sperm cysts can differ in how many nuclei are present at the beginning of meiosis.

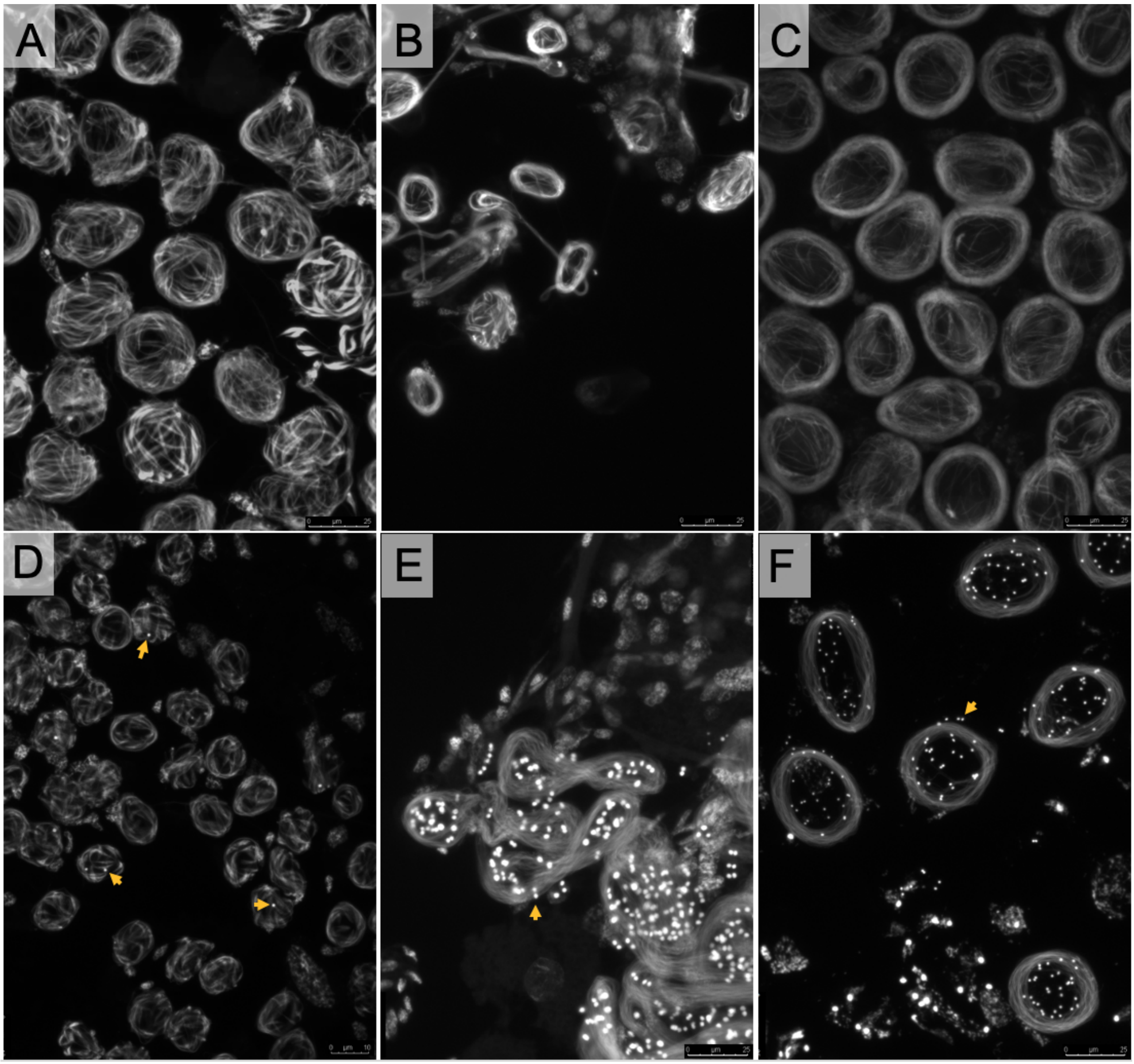


**Supplementary Figure 3**. Sperm bundles from (**A**) *Cystococcus campanidorsalis*, (**B**) *C. echiniformis*, (**C**) *C. pomiformis*, (**D**) *Ascelis praemollis*, (**E**) *Sphaerococcus ferrugineus*, and (**F**) *Eriococcus coriaceus*. Pycnotic nuclei are not present at the end of meiosis in males in (**A-C**), suggesting that the majority of paternal chromosomes do not take part in meiosis (i.e. are eliminated prior to meiosis). In *A. praemollis* (**D**), pycnotic nuclei (arrows) are present in a few, but not many sperm bundles (likely because most paternal chromosomes are eliminated before meiosis, with a small number participating in meiosis and becoming pycnotic nuclei afterwards). Both *S. ferrugineus* and *E. coriaceus* (**D-E**), do have pycnotic nuclei associated with sperm bundles.

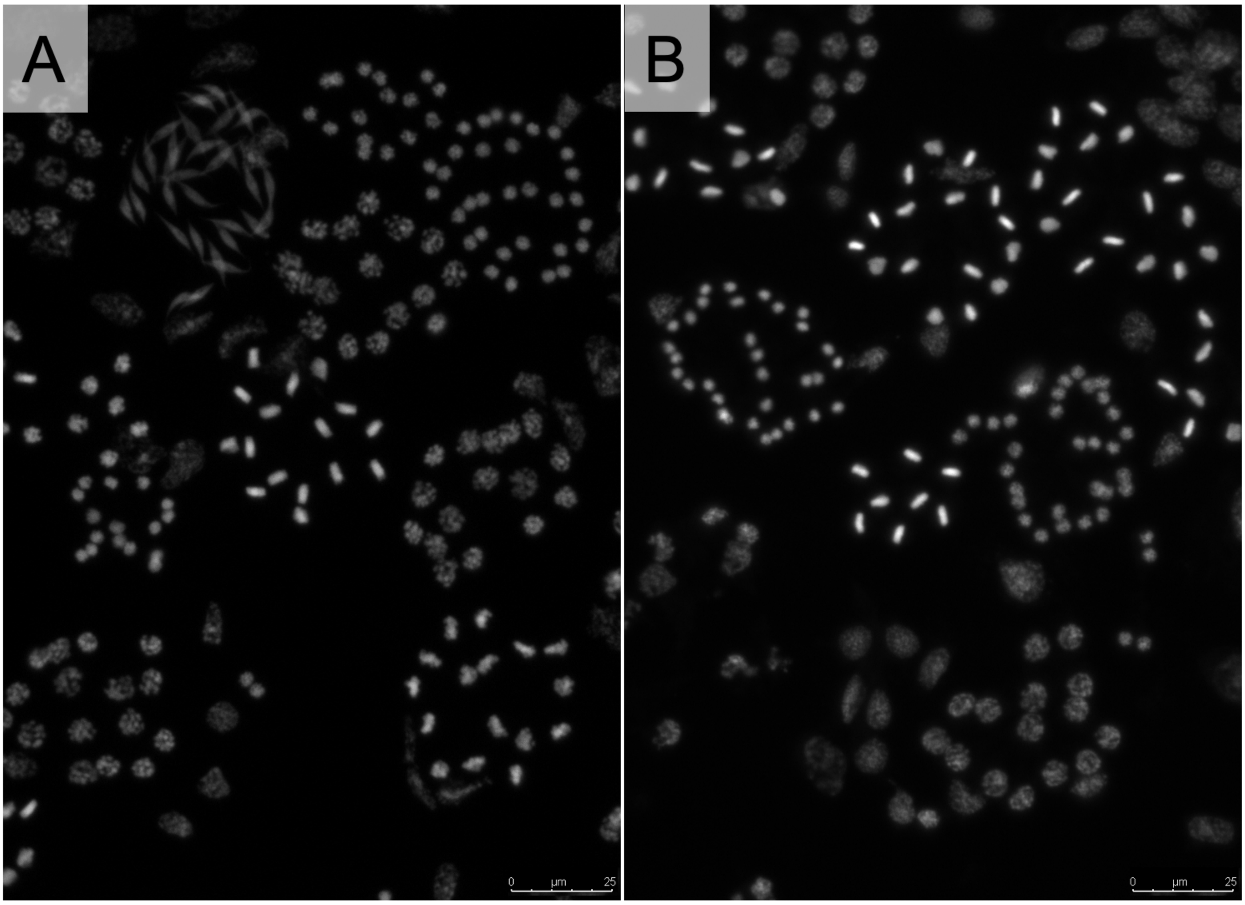


**Supplementary Figure 4.** Sperm cysts undergoing meiosis in *Cystococcus campanidorsalis* (**A**) and *C. echiniformis* (**B**). Sperm cysts in *C. echiniformis* can have either 8 or 16 primary spermatids in each sperm cyst, (both pictured) but cysts with 16 primary spermatids are much more common.

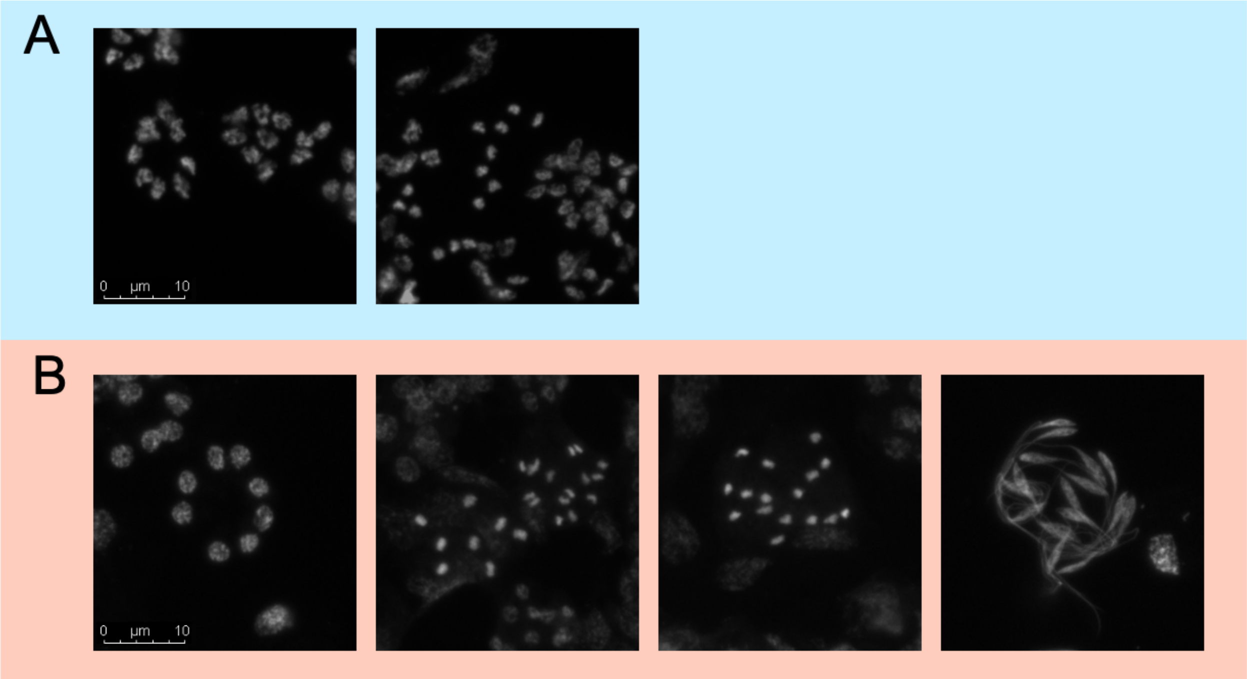


**Supplementary Figure 5**. Male meiosis in *Ascelis schraderi* (**A**) and *A. praemollis* (**B**). In both species male meiosis has one division with 8 nuclei in each sperm cyst at the beginning of meiosis and 16 sperm in each sperm bundle. This indicates that both species have Comstockiella PGE. Note that for *A. schraderi* we were not able to view sperm bundles forming and got information about number of sperm in sperm bundles from Brown (1967)

**Supplementary Table 5:** Model output analysing whether families inherit a different number of alleles depending on whether the allele is of maternal or paternal origin and whether the males are *C. echiniformis* or *C. campanidorsalis* males. The estimate and the standard error (in parentheses) is shown as well as whether the factor was predicted to be significant.

| | Number of alleles inherited |
|---|---|
| Species | 0.627(0.483) |
| Parent | 2.382 (0.503)*** |
| Constant | -0.342(0.438) |
| Observations | 38 |
| Log Likelihood | -65.297 |
| AIC | 138.595 |
| BIC | 145.145 |

Note: *p<0.05, **p<0.01, ***p<0.001

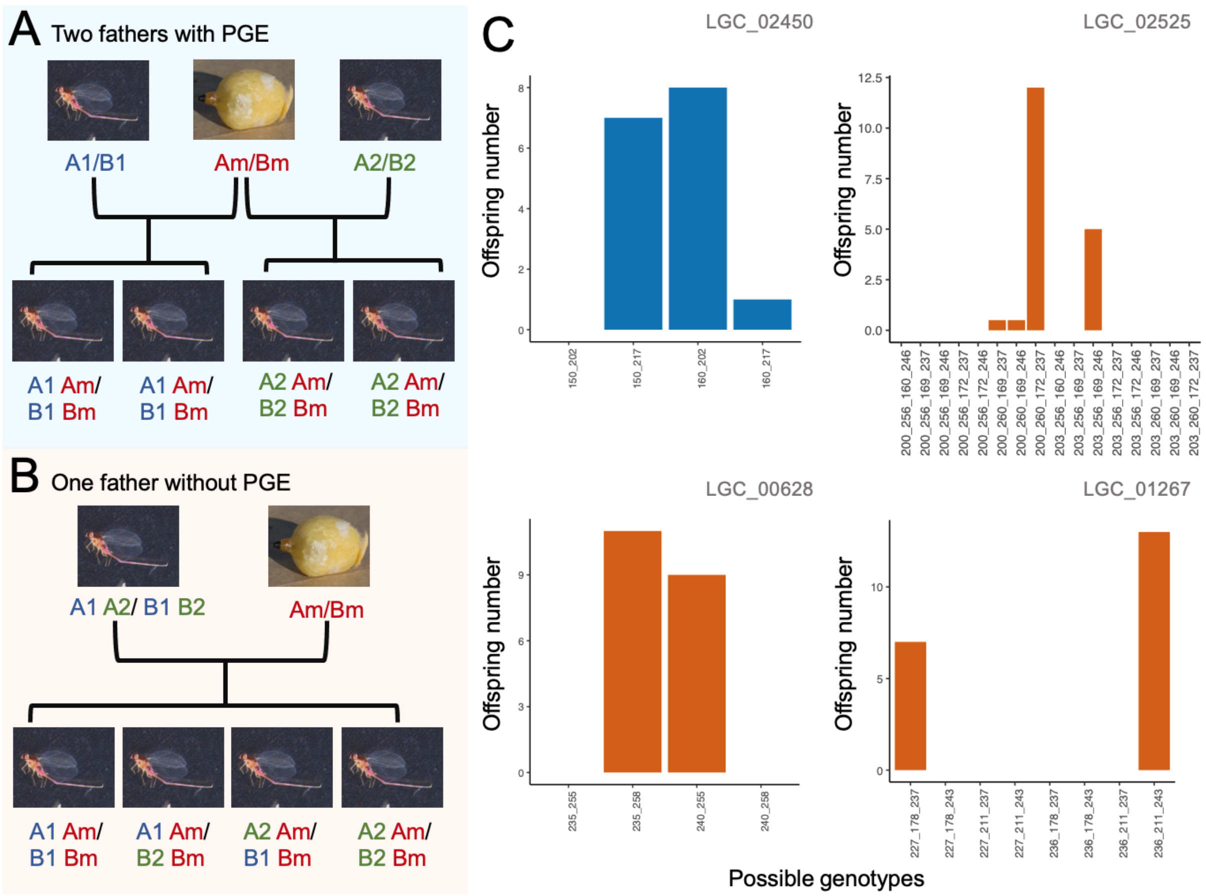


**Supplementary Figure 6.** (**A-B**) Schematic showing how *Cystococcus* families may have inherited more than one allele from their father. In (**A**) two fathers that exhibit PGE mated with the focal female (Am/Bm), producing two sons with the genotype of the first male, and two sons with the genotype of the second male (for both microsatellite loci A and B), while in (**B**), the female mated with one male who does not exhibit PGE and so passes the two alleles for A and B to his offspring randomly. The sons in this scenario are expected to have all possible combination of alleles. (**C**) Number of offspring with each allele combination in *Cystococcus campanidorsalis* (blue) and *C. echiniformis* (orange) families in which sons inherited two paternal alleles for two or more microsatellite loci. In all four families, not all allele combinations are present that would be expected if males had one father that was transmitting alleles in a Mendelian fashion. Rather, allele inheritance patterns are more similar to scenario (**A**). The values on the x-axis indicate the size of each PCR amplicon for each genotype.

**Supplementary Table 6**. In the four cases in which families inherited two paternal alleles for more than one microsatellite loci (one for *C. campanidorsalis*- LGC02450, and three for *C. echiniformis*), analysis of whether the inheritance patterns of alleles from the two or more sets of microsatellite loci were random. In all cases, Chi-square tests suggest that the allele inheritance is not random, likely because fathers of these families do exhibit PGE, but that the mothers of these families mated with more than one male. Family LGC_02450: 2 loci, LGC_00628: 2 loci, LGC_01267: 3 loci, LGC_02525: 4 loci.

| | LGC_02450 | LGC_00628 | LGC_01267 | LGC_02525 |
|---|---|---|---|---|
| Chi-square | 12.5 | 20.4 | 67.2 | 132.67 |
| d.f. | 3 | 3 | 7 | 15 |
| p-value | 0.00585 | 0.00014 | <0.0001 | <0.0001 |

**Supplementary Table 7:** Summary of number of homozygous (Hom) and heterozygous (Het) SNPs called for each sample in the RNAseq study analysing whether *C. campanidorsalis* and *C. echiniformis* males only express maternally inherited alleles. We performed Fishers exact tests comparing the number of homozygous and heterozygous alleles expressed in sons compared to their mother, although these tests generally suggest that the number of heterozygous SNPs was different between mothers and sons, both family members exhibited significant heterozygous expression of alleles for both species, with between 59-85% of alleles called as heterozygous.

| Species | Family | Individual | # SNP (after filtering) | # Hom SNP | # Het SNP | Total SNP in analysis | % Hom SNP | % Het SNP | Fishers test (p-value) | odds ratio |
|---|---|---|---|---|---|---|---|---|---|---|
| *C. echiniformis* | LGC_03572F4 | Female | 11803 | 3939 | 6870 | 10809 | 36.44% | 63.56% | | |
| | | Male1 | 14312 | 4792 | 8046 | 12838 | 37.33% | 62.67% | 0.0007333 | 1.114 |
| | | Male2 | 14202 | 4095 | 8073 | 12168 | 33.65% | 66.35% | 2.91E-10 | 1.201 |
| | LGC_03571F6 | Female | 10681 | 3596 | 6220 | 9816 | 36.63% | 63.37% | | |
| | | Male1 | 11624 | 4159 | 5989 | 10148 | 40.98% | 59.02% | 2.91E-10 | 1.201 |
| | | Male2 | 8155 | 2824 | 4383 | 7207 | 39.18% | 60.82% | 0.0007333 | 1.114 |
| | LGC_03571F5 | Female | 12679 | 4398 | 7244 | 11642 | 37.78% | 62.22% | | |
| | | Male1 | 15191 | 5398 | 8255 | 13653 | 39.54% | 60.46% | 0.004216 | 1.077 |
| | | Male2 | 13271 | 4523 | 7371 | 11894 | 38.03% | 61.97% | 0.6968 | 1.011 |
| *C. campanidorsalis* | LGC_03538F4 | Female | 25398 | 4961 | 17492 | 22453 | 22.10% | 77.90% | | |
| | | Male1 | 23272 | 3032 | 17847 | 20879 | 14.52% | 85.48% | 1.70E-05 | 0.889 |
| | | Male2 | 26841 | 3610 | 21007 | 24617 | 14.66% | 85.34% | 5.31E-05 | 0.900 |

**Supplementary Figure 7.** Image of *Ascelis praemollis* germ tissue, showing paternally inherited chromosomes potentially being eliminated from cells (arrows). As these cells have not yet undergone meiosis, this suggests that chromosome elimination prior to meiosis in this species may be common. However, this image also suggests that different numbers of chromosomes may be eliminated prior to meiosis, as the size of the chromosomes being eliminated varies for different cells.

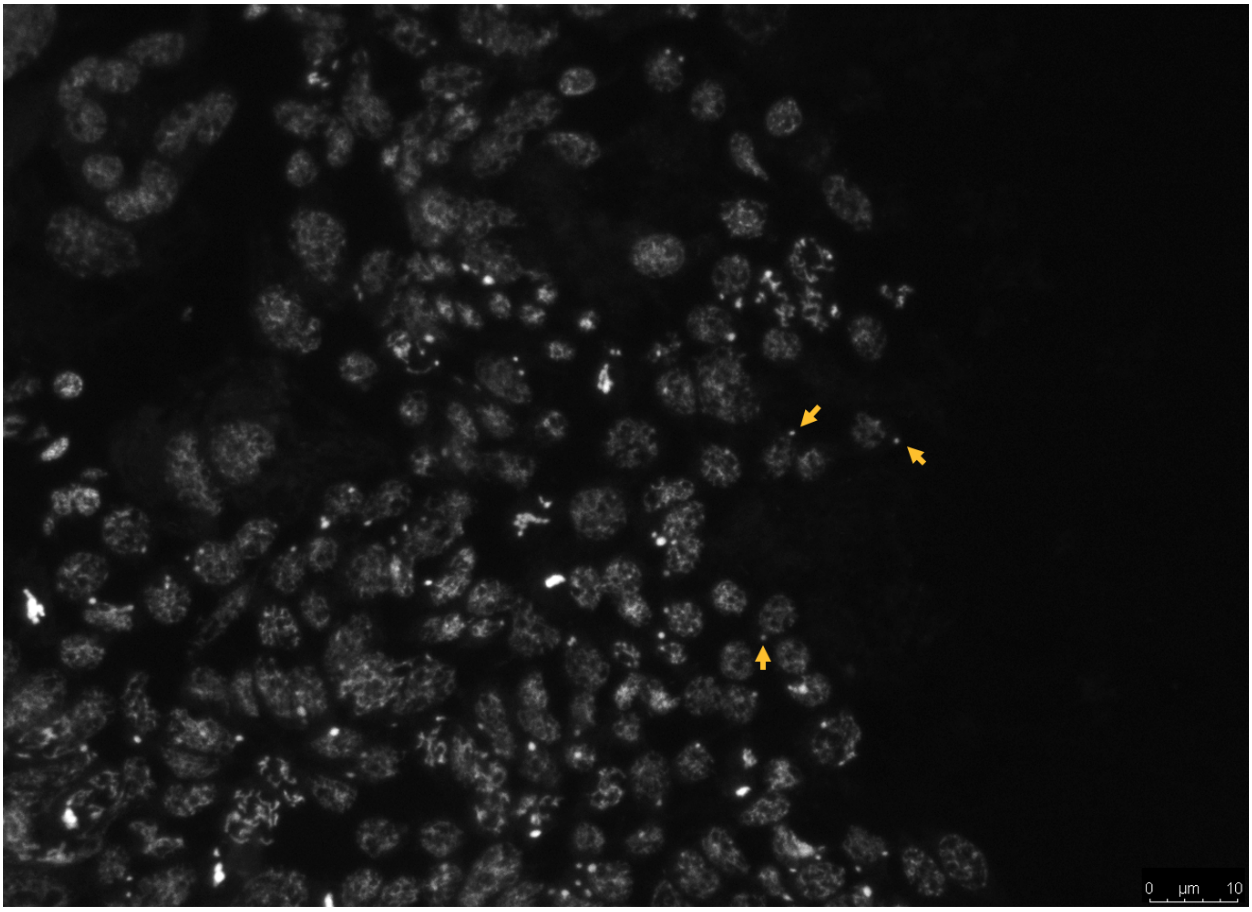
